# PRDX1 regulates T cell effector function in the ovarian tumor microenvironment

**DOI:** 10.64898/2026.08.21.746361

**Authors:** Sarah J. McPhedran, Gillian Carleton, Scott Hannan, Sarah MacPherson, Leonardo Castro, Sam Preshaw, Julian J. Lum

## Abstract

T cell-based immunotherapies have remained ineffective against high-grade serous ovarian carcinoma (HGSOC). The metabolic environment of HGSOC suppresses the activity of cellular therapies, however, the metabolites that enhance or suppress T cell antitumor activity are not fully understood. Here, a pooled CRISPR-Cas9 knockout screen in primary human T cells cultured with patient-derived ascites was used to identify metabolic enzymes that inhibit effector cytokine production and cytolytic function. The screen identified PRDX1 as a negative regulator of T cell effector function. Targeted deletion of *PRDX1* increased the frequency of IFN-γ-producing T cells, enhanced glucose uptake, increased mitochondrial mass, and improved T cell viability under suppressive ascites conditions. Mechanistically, PRDX1 deficiency increased intracellular reactive oxygen species (ROS) and impaired autophagic flux. The effects of *PRDX1* deletion enhanced aspects of T cell function, while its effects on chimeric antigen receptor (CAR)-T cell cytotoxicity were donor dependent. Collectively, this study identifies PRDX1 as a regulator of T cell activation, metabolism, and effector function in the inhibitory physiological suppressive environment of HGSOC ascites.

## Introduction

T cell-based immunotherapies have transformed the treatment landscape for numerous cancers, including leukemia, lymphoma, multiple myeloma, and melanoma (*1*, *2*). Although these therapies have only recently entered clinical practice, their rapid development and expanding applications suggest that their impact on cancer treatment is only beginning to be realized (*3*, *4*). Continued research is therefore required to fully harness the therapeutic potential of T cell-based immunotherapy, particularly in solid tumors.

High-grade serous ovarian carcinoma (HGSOC) is one example of a solid tumor that has shown limited responsiveness to immunotherapies, including chimeric antigen receptor (CAR) T cell therapy (*5*, *6*). Multiple barriers contribute to this lack of efficacy in solid tumors, including poor T cell infiltration, low neoantigen burden, and a highly immunosuppressive tumor microenvironment (TME) (*5*). While the TME suppresses antitumor immunity through numerous mechanisms, including inhibitory cytokines and suppressive immune cell populations, metabolites in the TME have emerged as a major obstacle to effective T cell function in HGSOC (*7–9*).

Metabolic suppression within the TME can broadly be categorized into two major mechanisms. First, tumor cells consume large quantities of essential nutrients to support uncontrolled growth, resulting in T cell metabolic bankruptcy: the depletion of metabolites such as glucose, glutamine, and tryptophan from the surrounding environment (*10*, *11*). In contrast, T cells, which are primed in nutrient-replete environments such as the thymus and lymphoid organs, are poorly adapted to compete for these scarce resources (*12*). Second, tumor, stromal and anti-inflammatory immune cells (e.g. myeloid-derived suppressor cells [MDSCs]) produce and secrete immunosuppressive metabolites that directly induce T cell metabolic arrest causing impaired function as a result of their dysregulated metabolism (*13*). These metabolites include kynurenine, methylglyoxal, and lactate, which accumulate in the TME and contribute to T cell dysfunction (*13*).

To overcome these barriers, the field of metabolic engineering seeks to modify T cells to enhance their metabolic fitness and functionality within metabolically hostile tumor environments (*14*). Several metabolic genes have been identified in preclinical studies, and a growing number are being evaluated in clinical trials, such as Regnase-1 and the A2A receptor (*15–20*). However, our understanding of the metabolic pathways that negatively regulate T cell function in the TME remains incomplete.

In this study, the identity of metabolic genes that act as negative regulators of T cell effector function in the suppressive HGSOC ascites microenvironment was investigated. HGSOC has been shown to be enriched in multiple immunosuppressive proteins and metabolites, thereby providing an environment that recapitulates key features of the TME (*9*, *21*). An optimized pooled CRISPR-Cas9-knockout screening platform was developed using primary human T cells cultured in patient HGSOC ascites as a physiologically relevant model that represents critical metabolic aspects of the TME. This screen identified *PRDX1* as a negative regulator of T cell function, demonstrating that *PRDX1* deletion enhances T cell effector function under suppressive ascites conditions. Further characterization revealed that *PRDX1*-knockout T cells exhibit enhanced activation, altered metabolic phenotypes, and donor-dependent improvements in CAR-T cell cytotoxicity. *In vitro, PRDX1* regulates T cell function through a mechanism involving reactive oxygen species (ROS) homeostasis and autophagy regulation. Collectively, this work highlights PRDX1 as a potential target for metabolic engineering of therapeutic T cells.

## Results

### HGSOC ascites supernatant recapitulates key immunosuppressive features of the tumor microenvironment

To identify metabolic regulators of T cell function in high-grade serous ovarian carcinoma (HGSOC), an ex vivo model was established that recapitulates key features of the ovarian cancer tumor microenvironment (TME). Malignant ascites is a hallmark of advanced HGSOC and contains tumor cells, stromal cells, immune cells, soluble factors, and metabolites that collectively contribute to immune suppression (Fig. 1A) (*22*, *23*). We therefore hypothesized that patient ascites supernatant could serve as a physiologically relevant selection pressure for functional genetic screening.

**Fig. 1.**
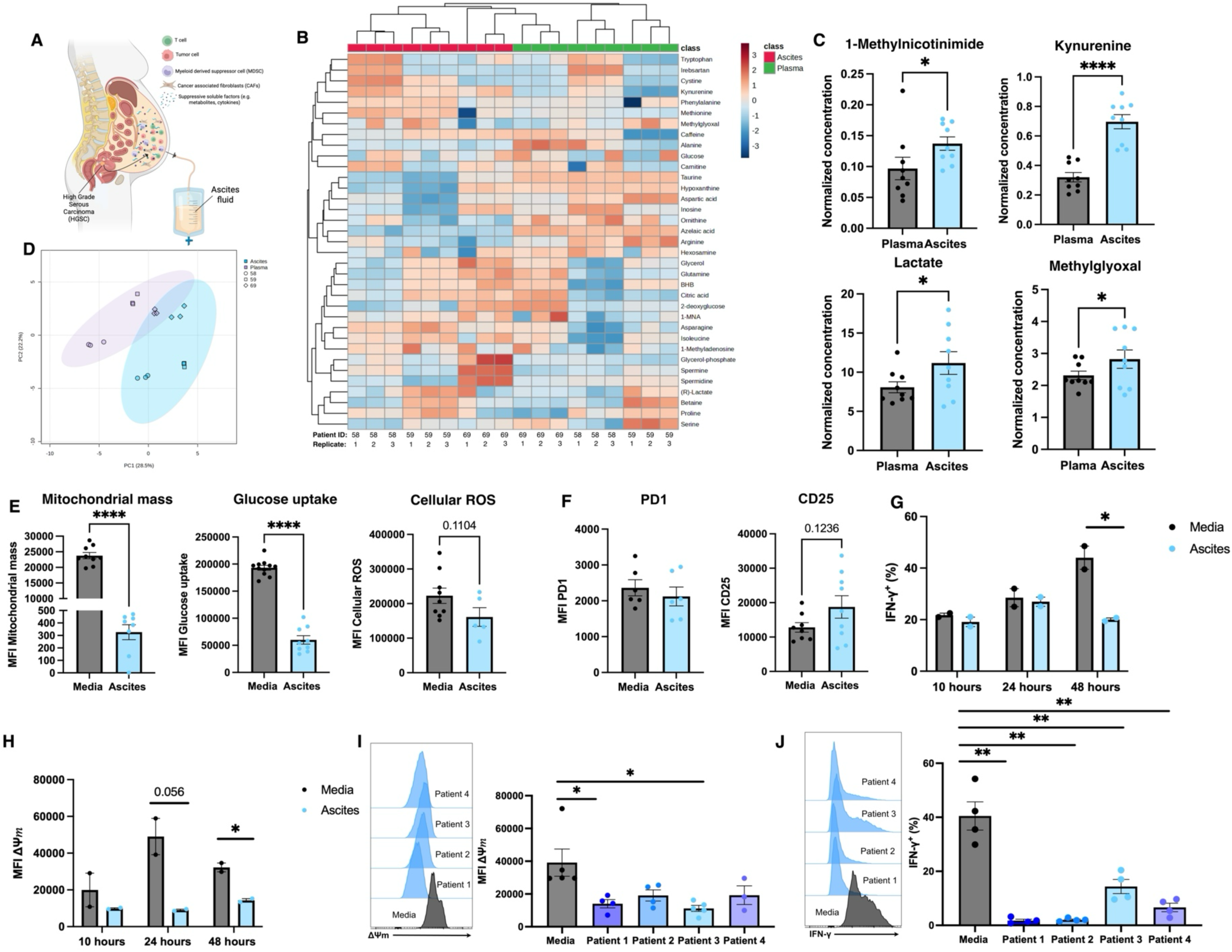
HGSOC ascites supernatant recapitulates a metabolically suppressive tumor microenvironment that impairs T cell function. **(A)** Schematic illustrating collection of ascites fluid from patients with high-grade serous ovarian carcinoma (HGSOC) and its use as an ex vivo model of the ovarian tumor microenvironment (TME). Created with BioRender. **(B)** Heatmap showing the relative abundance of pre-selected metabolites detected by untargeted metabolomic profiling of patient-matched plasma and ascites samples from three patients with HGSOC. Peak intensities were filtered by interquartile range, normalized by sum, log-transformed, and auto-scaled prior to analysis. Samples are displayed in columns and metabolites in rows. Hierarchical clustering was performed using Euclidean distance and average linkage. Heatmap was generated using MetaboAnalyst 6.0. **(C)** Selected metabolites significantly enriched in ascites relative to matched plasma, including 1-methylnicotinamide, kynurenine, lactate, and methylglyoxyl. Each point represents a biological or technical replicate (3 different patients, with each patient sample collected in triplicate). **(D)** Principal component analysis (PCA) of metabolomic peak intensities (using pre-selected metabolites from heatmap) from patient-matched plasma and ascites samples following interquartile range filtering, normalization by sum, log transformation, and auto-scaling. Patient identity was included as metadata. The PCA demonstrates distinct metabolic profiles between plasma and ascites samples and group separation was statistically significant by PERMANOVA (P = 0.001). **(E)** Metabolic characterization of primary human CD3+ T cells following 48 hr stimulation cultured in either complete media or 100% patient ascites. Data are representative of a single T cell and ascites donor. Each point represents a technical replicate. **(F)** Expression of activation and exhaustion-associated markers on CD3+ T cells following 48 hr stimulation in media or 100% ascites. Data are representative of a single T cell and ascites donor. Each point represents a technical replicate. **(G)** Intracellular IFN-γ production by CD3+ T cells stimulated and cultured in media or ascites for 10, 24, or 48 hrs. Data are representative of a single T cell and ascites donor. Each point represents a technical replicate. Statistical significance was determined using unpaired Welch’s t-tests comparing media and ascites conditions at each timepoint. **(H)** Mitochondrial membrane potential (ΔΨm) of stimulated CD3+ T cells cultured in media or ascites for 10, 24, or 48 hrs. Data are representative of a single T cell and ascites donor. Each point represents a technical replicate. Statistical significance was determined using unpaired Welch’s t-tests comparing media and ascites conditions at each timepoint. **(I)** Representative histograms and quantification of mitochondrial membrane potential in CD3+ T cells cultured for 48 hrs in media or ascites obtained from four independent HGSOC patients. Data were generated using two independent T cell donors. Each point represents a biological or technical replicate. Media and ascites conditions were compared independently for each ascites sample using Welch’s t-test. **(J)** Representative histograms and quantification of IFN-γ production in CD3+ T cells cultured for 48 hrs in media or ascites obtained from four independent HGSOC patients. Data were generated using two independent T cell donors. Each point represents a biological or technical replicate. Media and ascites conditions were compared independently for each ascites sample using Welch’s t-test. For C, E-J: Bars represent mean ± SEM; statistical significance was determined using Welch’s t-test, *P < 0.05, **P < 0.01, ***P < 0.001, ****P < 0.0001.

To determine whether ascites possesses a distinct metabolite composition compared with plasma, we performed metabolomic profiling of patient-matched plasma and ascites supernatant samples from patients with HGSOC. Patient-matched plasma and ascites samples exhibited distinct metabolomic abundances (Fig. 1B). Specifically, several metabolites previously implicated in T cell suppression were enriched in ascites, including kynurenine, lactate, 1-methylnicotinamide and methylglyoxal (Fig. 1C) (*9*, *24*–*26*). Despite the enrichment of several immunosuppressive metabolites, nutrients known to be depleted in HGSOC tumors, such as glucose and glutamine, were not reduced in ascites compared with plasma, with glutamine levels instead being elevated (Fig. 1B) (*27*). This observation suggests that the HGSOC tumor microenvironment may be less metabolically bankrupt but instead be enriched for suppressive metabolites that induce T cell arrest. Principal component analysis (PCA) of pre-selected metabolites known to be important for T cell function further demonstrated distinct clustering of plasma and ascites specimens, indicating that ascites represents a unique physiological medium (Fig. 1D). Together, these findings demonstrate that HGSOC ascites possesses a metabolically distinct composition enriched for factors known to impair T cell function.

Next, the functional impact of ascites on primary human T cells was assessed. T cells were activated in the presence of either standard culture media or ascites supernatant and analyzed for metabolic, effector and activation phenotypes. Exposure to ascites resulted in a greater than 75-fold reduction in mitochondrial mass and glucose uptake compared with media controls (Fig. 1E), consistent with a metabolically suppressive environment. Cellular reactive oxygen species (ROS) levels also trended lower in ascites-treated cells, although this difference did not reach statistical significance (Fig. 1E). While expression of the activation marker PD-1 remained unchanged, ascites-exposed T cells exhibited a trend toward increased CD25 expression, suggesting that impaired activation is unlikely to be a dominant force of the suppressive effects of ascites (Fig. 1F).

Furthermore, ascites significantly reduced IFN-γ production following T cell activation, with suppression becoming most apparent after 48 hours of culture (Fig. 1G). Similarly, mitochondrial membrane potential (ΔΨm), measured by MitoTracker Deep Red staining, was reduced in ascites-treated T cells at multiple time points (Fig. 1H). To evaluate whether T cell suppression was consistent across multiple patient ascites samples, T cells were cultured in ascites obtained from four independent HGSOC donors. All ascites samples significantly reduced mitochondrial membrane potential and IFN-γ relative to media controls, although the magnitude of suppression varied between donors (Fig. 1I-J).

Collectively, these data demonstrate that HGSOC ascites recapitulates key immunosuppressive metabolic features of the ovarian cancer TME and consistently suppresses T cell mitochondrial activity and effector function. These findings establish HGSOC patient ascites supernatant as a biologically relevant selection pressure for identifying metabolic regulators of T cell effector function using pooled CRISPR-Cas9 screening. Because mitochondrial membrane potential and interferon gamma (IFN-γ) production were dynamically suppressed by ascites, these phenotypes were selected as the primary readouts of T cell effector function in our CRISPR screen.

### Development and optimization of a pooled CRISPR-Cas9 screen to identify metabolic regulators of T cell function in HGSOC ascites

Having established patient ascites supernatant as a metabolically suppressive model of the HGSOC tumor microenvironment, a pooled CRISPR-Cas9-knockout screening platform was developed to identify metabolic genes that regulate IFN-γ and mitochondrial membrane potential under these conditions. The CRISPR-Cas9 screening method was adapted from the single guide RNA (sgRNA) lentiviral infection with Cas9 protein electroporation (SLICE) method, outlined in Shifrut et al (*28*). The complete screening workflow is shown in Fig. 2A.

**Fig. 2.**
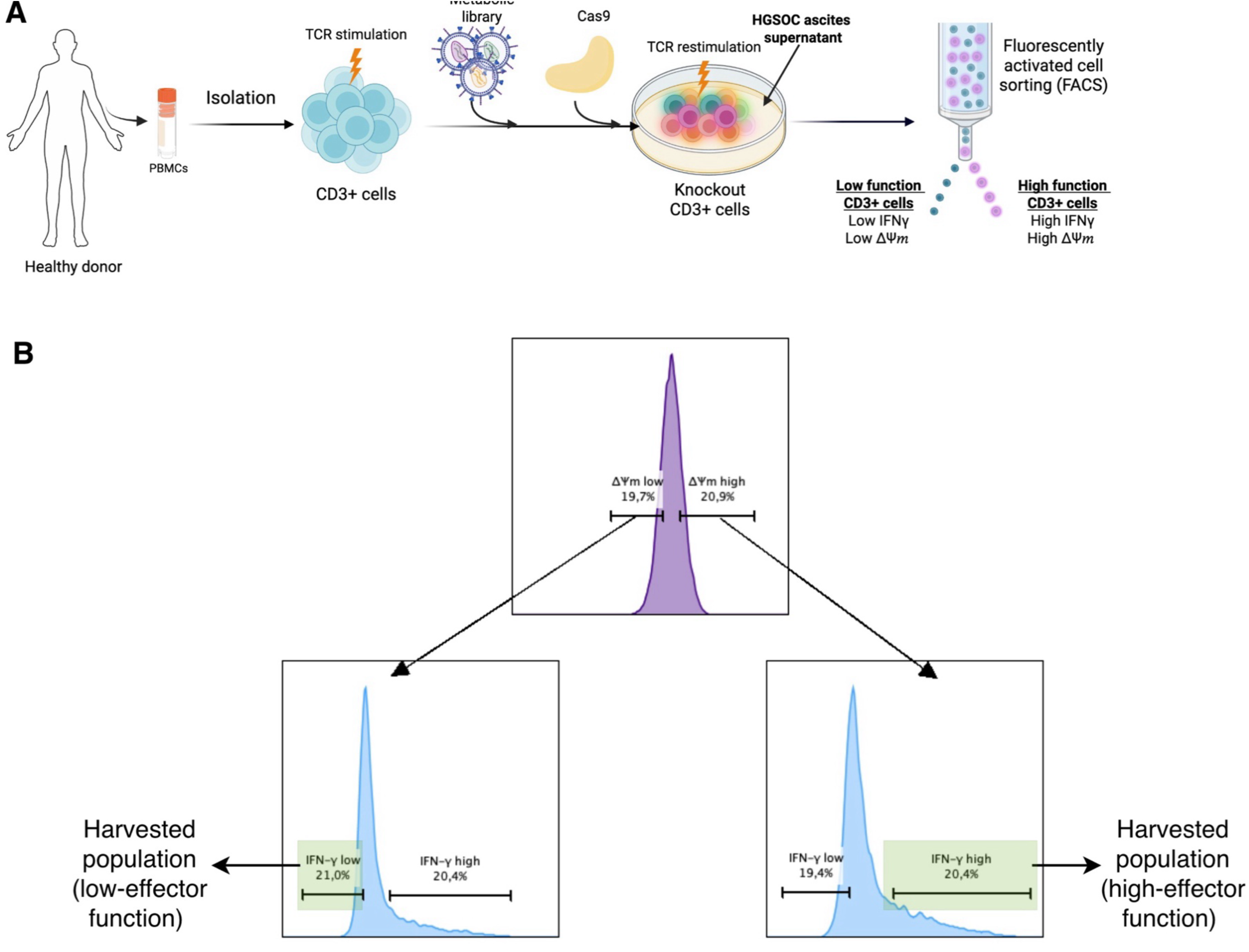
Pooled CRISPR-Cas9 screening strategy to identify metabolic gene knockouts that enhance T cell function in suppressive HGSOC ascites. **(A)** Schematic of the pooled CRISPR-Cas9 screening workflow. CD3+ T cells were transduced with a metabolism-focused sgRNA library (29,903 sgRNAs targeting 2,981 genes) and electroporated with Cas9. Edited cells were cultured in HGSOC ascites, restimulated for 48 hrs, and FACS-sorted into high- and low-effector populations based on IFN-γ production and mitochondrial membrane potential (ΔΨm). sgRNAs were then quantified by next-generation sequencing. **(B)** FACS gating strategy used to isolate high-effector (IFN-γ-high; ΔΨm-high) and low-effector (IFN-γ-low; ΔΨm-low) T cell populations for sequencing. Sequencing data were analyzed using MAGeCK to identify enriched metabolic gene knockouts.

Based on the initial ascites assays above (Fig. 1G-J), IFN-γ and mitochondrial membrane potential (ΔΨm) were selected as orthogonal readouts of T cell effector function and metabolic fitness. Cells exhibiting high IFN-γ and high ΔΨm were classified as high-effector function, whereas cells exhibiting low IFN-γ and low ΔΨm were classified as low-effector function (Fig. 2A,B). Genomic DNA was subsequently isolated from these two sorted populations for sgRNA quantification by next-generation sequencing.

The kinetics of IFN-γ production following T cell receptor (TCR) restimulation was optimized for use in the screen. Although IFN-γ-positive cells were detectable after 10 hours of stimulation, a larger and more reproducible IFN-γ-positive population was observed following 48 hours of restimulation across independent experiments (Fig. S1A). Therefore, a 48-hour restimulation period was selected for all subsequent screening experiments to maximize both the magnitude and reproducibility of the IFN-γ response.

Next, the generation and expansion of pooled-knockout T cells following lentiviral transduction with the metabolic sgRNA library and Cas9 electroporation was assessed. Puromycin selection successfully enriched for sgRNA-containing cells, resulting in robust expansion of the library-transduced population, whereas non-transduced cells failed to proliferate under puromycin selection conditions (Fig. S1B). Having established efficient library generation, flow cytometric sorting was used to isolate functionally distinct T cell populations. Cells were gated on mitochondrial membrane potential (ΔΨm) and IFN-γ expression, and the upper and lower 20% of each distribution were harvested to generate IFN-γ/ΔΨm-high and IFN-γ/ΔΨm-low populations, for downstream sequencing analysis (Fig. 2B; Fig. S1C).

To ensure adequate representation of the metabolic sgRNA library throughout the screen, coverage calculations using the final library size of 29,903 sgRNAs were performed. These calculations aimed for a 100-fold representation per sgRNA, as outlined in Bock et al (*29*), and anticipated sorting frequencies of 20% for both ΔΨm and IFN-γ gates (Fig. 2B; Fig. S1C) and assumed 3-fold expansion of pooled-knockout cells (Fig. S1B). We determined that approximately 5.6×10^7^ starting T cells would be required to maintain sufficient library coverage through expansion and sorting (Fig. S1D). Under these conditions, each harvested population exceeded the desired 100-fold sgRNA representation threshold.

Finally, the effects of lentiviral transduction, puromycin selection, and Cas9-mediated genome editing on the phenotypes used as screen readouts (ΔΨm and IFN-γ) were assessed in the pooled knockout population. Comparison of library-transduced cells (no Cas9) and pooled knockout cells (library and Cas9) demonstrated a significant reduction in IFN-γ production following genome editing, whereas mitochondrial membrane potential remained unchanged (Fig. S1E). The observed reduction in IFN-γ production is consistent with successful pooled gene disruption, implying that there was efficient editing with the library-encoded targets. Importantly, both readouts retained sufficient dynamic range to support phenotypic segregation by flow cytometry.

Collectively, these experiments established a robust pooled CRISPR-Cas9 screening platform capable of maintaining library coverage in primary human T cells while enabling functional selection based on IFN-γ production and mitochondrial activity under inhibitory HGSOC ascites conditions.

### Pooled CRISPR-Cas9 screen identifies PRDX1 as a negative regulator of T cell function in HGSOC ascites

Following screening optimization, a pooled CRISPR-Cas9-knockout screen was performed to identify metabolic genes that regulate T cell function under suppressive ascites from patient 3 (see Fig. 1I-J). Primary human T cells transduced with the metabolic sgRNA library were cultured in ascites and sorted into IFN-γ/ΔΨm-high and IFN-γ/ΔΨm-low populations prior to next-generation sequencing of sgRNA abundance (Fig. 2A).

To assess screen quality, sequencing and library representation metrics were first examined. Across all sorted populations, 76–80% of sequencing reads successfully mapped to library sgRNAs (Fig. S2A). Analysis of sgRNA distribution revealed low Gini indices within the starting library population, indicating relatively uniform sgRNA representation prior to selection (Fig. S2B). Principal component analysis of MAGeCK-normalized sgRNA counts demonstrated clear separation between the IFN-γ/ΔΨm-high and IFN-γ/ΔΨm-low populations, while the library sequences clustered independently (Fig. S2C), suggesting that ascites-mediated selection generated distinct functional T cell populations.

To identify candidate regulators of T cell function, sgRNA abundance was compared between IFN-γ/ΔΨm-high and IFN-γ/ΔΨm-low populations using MAGeCK analysis (*30*). Numerous genes were significantly enriched or depleted in the high-effector function population, represented in blue or pink, respectively (Fig. 3A). Genes enriched within the IFN-γ/ΔΨm-high population represent candidate negative regulators of T cell function, as their disruption enhanced T cell fitness under inhibitory ascites conditions. Conversely, genes depleted within the IFN-γ/ΔΨm-high population represent candidate positive regulators of T cell function. Notably, *NFKB1*, which encodes a subunit of the well-established pro-inflammatory transcription factor NF-κB that promotes T cell effector function (*31*), was depleted in the high-population, providing biological support for the validity of the screen.

**Fig. 3.**
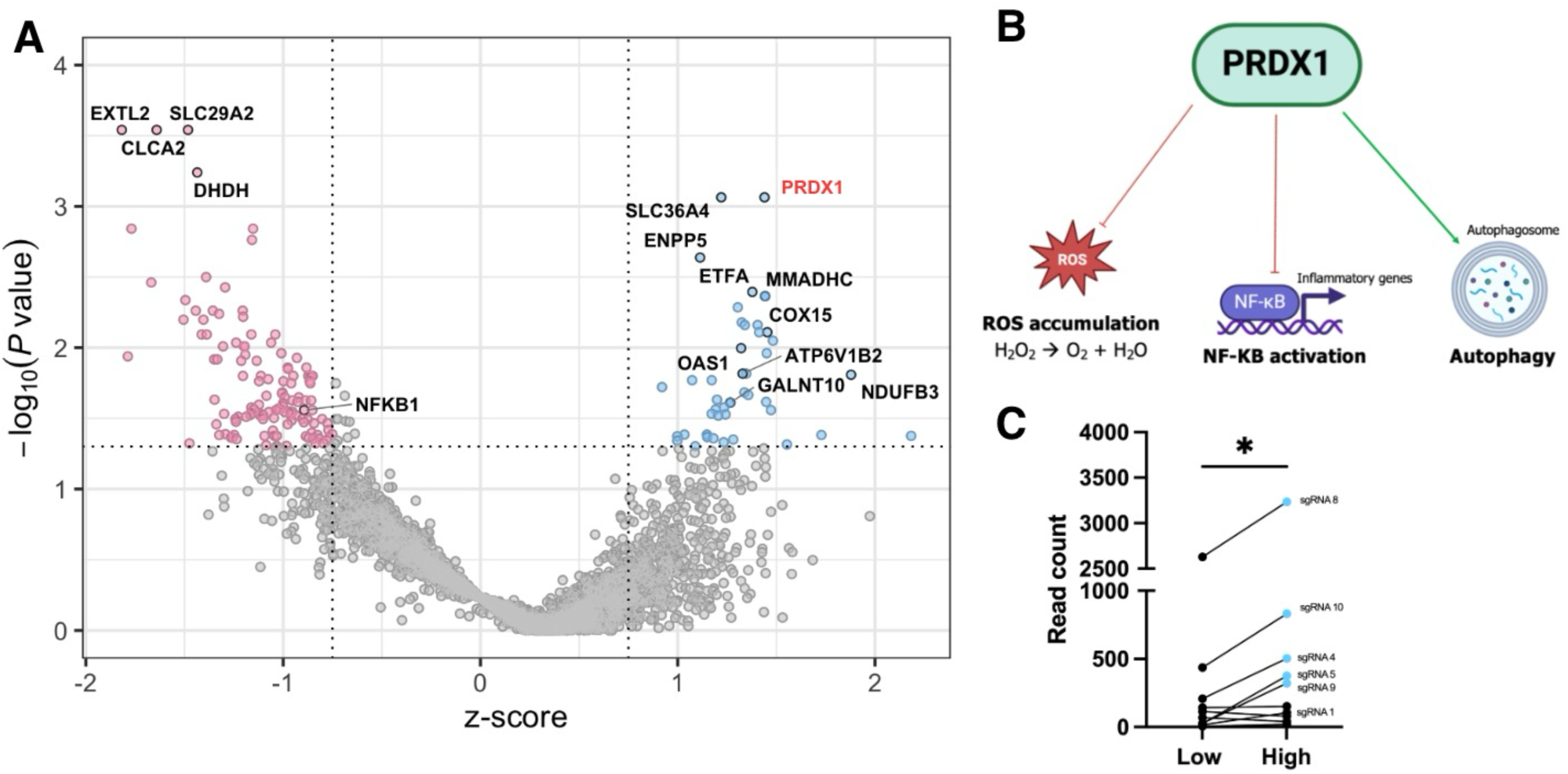
PRDX1 is identified as a candidate negative regulator of T cell effector function in suppressive ascites. **(A)** Volcano plot displaying MAGeCK maximum-likelihood estimation (MLE) analysis of the pooled CRISPR-Cas9 screen performed in primary human T cells cultured in suppressive ovarian cancer ascites (Fig. 1I-J: patient 3). Each point represents a single gene targeted in the screen (2,981 genes total). Positive Z-scores indicate enrichment of gene knockouts in the high-effector function population (IFN-γ-high; ΔΨm-high) and negative Z-scores indicate depletion from the high-effector function population. **(B)** Known pathways regulated by PRDX1 include ROS homeostasis, NF-κB activation, and promoting autophagy induction. Created with BioRender. **(C)** Read counts for individual *PRDX1*-targeting sgRNAs in low-effector function (IFN-γ-low; ΔΨm-low) and high-effector function (IFN-γ-high; ΔΨm-high) sorted populations. Statistical significance was determined using a paired t-test. *P < 0.05.

Among all candidate genes, *PRDX1* emerged as the most significant hit enriched within the IFN-γ/ΔΨm-high population (Fig. 3A). *PRDX1* encodes peroxiredoxin-1, an antioxidant enzyme involved in regulating cellular reactive oxygen species (ROS) homeostasis that has also been implicated in NF-κB signaling and autophagy pathways (Fig. 3B) (*32–35*). Examination of individual sgRNA representation revealed enrichment of five independent *PRDX1*-targeting sgRNAs within the high-effector function population relative to the low-effector function population (Fig. 3C), supporting an on-target effect rather than enrichment driven by a single sgRNA.

Based on its strong statistical enrichment, consistent sgRNA behavior, and established roles in cellular pathways linked to metabolism and function, *PRDX1* was selected for downstream validation studies.

### PRDX1-knockout increases the frequency of IFN-γ-producing T cells but does not alter mitochondrial membrane potential

To validate *PRDX1* as a regulator of T cell function and mitochondrial activity, we generated *PRDX1*-knockout primary human T cells using Cas9 ribonucleoprotein (RNP) electroporation and evaluated their phenotype (IFN-γ and ΔΨm) following culture in suppressive HGSOC ascites or standard media conditions (Fig. S3A). Three independent sgRNAs targeting *PRDX1* were selected based on their enrichment in the pooled CRISPR screen (Fig. 3C). TIDE analysis demonstrated efficient genome editing for all three sgRNAs, while western blot analysis confirmed loss of PRDX1 protein expression following gene deletion, particularly for sgRNAs 1 and 10 (Fig. 4A-B).

**Fig. 4.**
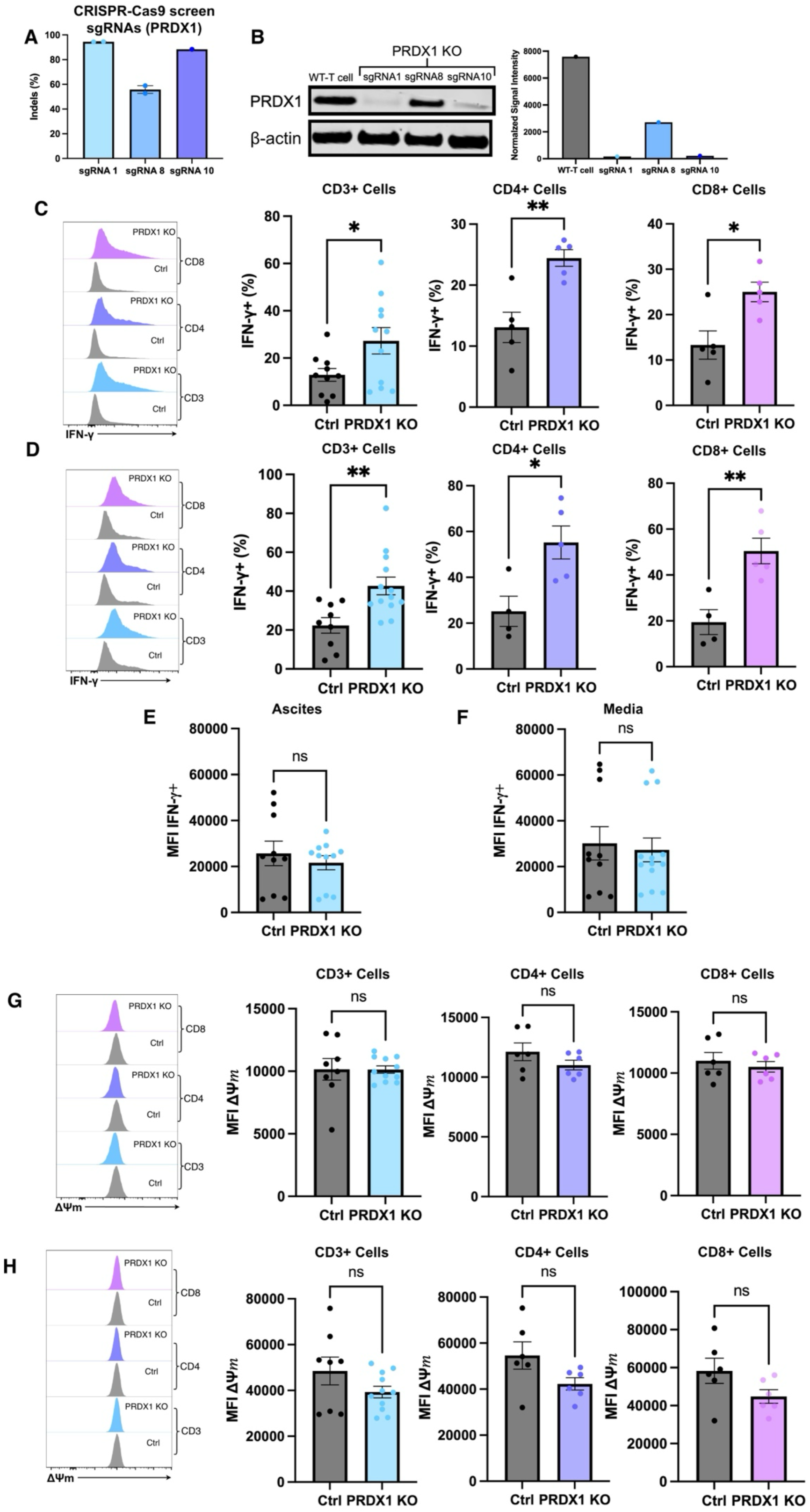
PRDX1 deletion increases IFN-γ-producing cells but does not alter mitochondrial membrane potential. **(A)** Indel generation of *PRDX1*-targeting sgRNAs identified in the CRISPR screen. Indel frequencies were determined by TIDE analysis of genomic DNA harvested from expanded T cells 14 days after isolation and activation and 11 days following Cas9 ribonucleoprotein (RNP) electroporation. Data are representative of one T cell donor. **(B)** Representative western blot showing PRDX1 expression in control T cells and T cells edited with *PRDX1*-targeting sgRNAs (sgRNA 1, sgRNA 8, and sgRNA 10 from Fig. 3B). Lysates were harvested 14 days after T cell isolation and activation and 11 days following RNP electroporation. Densitometric quantification of PRDX1 protein expression was normalized to β-actin. **(C–D)** Representative histograms and quantification are shown for the total CD3+ T cell population and the CD4+ and CD8+ T cell subsets. Intracellular IFN-γ production following *PRDX1* deletion. Control T cells and PRDX1-knockout T cells were stimulated for 48 hrs in either ascites **(C)** or media **(D). (E–F)** Mean fluorescence intensity (MFI) of IFN-γ-positive CD3+ T cells following 48 hr stimulation in ascites **(E)** or media **(F). (G– H)** Representative histograms and quantification are shown for the total CD3+ T cell population and the CD4+ and CD8+ T cell subsets. Mitochondrial membrane potential (ΔΨm) following 48 hr stimulation in ascites **(G)** or media **(H)**. For panels (C–H), T cells were cultured in media or ascites derived from the same HGSOC patient sample used in the pooled CRISPR screen (Fig. 1I-J: patient 3). Control T cells represent cells treated with no sgRNA, intergenic sgRNA, or *AAVS1*-targeting sgRNA; *PRDX1*-knockout T cells represent cells treated with sgRNA 1, sgRNA 8, and sgRNA 10 (from Fig. 3B). Each point represents a distinct sgRNA tested in a single T cell donor; some sgRNAs were evaluated in technical replicates across multiple experiments. Bars represent mean ± SEM. Statistical significance was determined using Welch’s t-test. *P < 0.05, **P < 0.01; ns, not significant.

PRDX1 expression was also assessed at baseline, throughout T cell culture, and activation to determine whether its expression was maintained over time. PRDX1 mRNA expression increased significantly between day 7 and day 14 of T cell expansion (Fig. S3D), indicating that PRDX1 is upregulated either during prolonged ex vivo culture or reactivation (or both). Following stimulation in ascites supernatant, PRDX1 expression remained lower than that observed in media but was not significantly different from day 7. As expected, PRDX1 transcript abundance was markedly absent in *PRDX1*-knockout T cells treated with sgRNA 10 (Fig. S3D).

We then sought to determine whether deletion of *PRDX1* recapitulated the phenotype observed in the pooled screen (Fig. 3A). *PRDX1*-knockout significantly increased the frequency of IFN-γ-positive cells within total CD3+, CD4+, and CD8+ T cell populations under both ascites and media conditions (Fig. 4C-D). To elucidate sgRNA-specific effects, results were stratified by control conditions (no sgRNA, intergenic sgRNA and AAVS1 sgRNA) and *PRDX1* targeting sgRNAs (sgRNA 1, 8 and 10). Under ascites conditions and media conditions, all three *PRDX1*-targeting sgRNAs increased the frequency of IFN-γ-producing T cells relative to control populations, with sgRNAs 8 and 10 producing the largest effects (Fig. S3B). Notably, the increase in IFN-γ-producing cells upon *PRDX1*-knockout was not accompanied by changes in IFN-γ mean fluorescence intensity (MFI) among IFN-γ positive cells (Fig. 4E-F), suggesting that *PRDX1*-knockout increases the proportion of T cells capable of producing IFN-γ rather than the amount of cytokine produced on a per-cell basis.

In contrast to the pooled screen results, *PRDX1*-knockout had minimal effects on mitochondrial membrane potential (ΔΨm) under either ascites or media conditions, suggesting that enrichment of *PRDX1*-targeting sgRNAs in the IFN-γ/ΔΨm-high population was driven primarily by enhanced IFN-γ production rather than increased mitochondrial activity (Fig. 4G-H; Fig. S3C).

Beyond the screening T cell donor, the editing efficiency of the three PRDX1-targeting sgRNAs (sgRNAs 1, 8, and 10) was evaluated across multiple healthy donors. Editing efficiencies varied between donors and sgRNAs, with sgRNA 8 exhibiting greater donor-to-donor variability than sgRNAs 1 and 10 (Fig. S3E). Furthermore, to assess the reproducibility of the increased frequency of IFN-γ-producing T cells following *PRDX1* deletion across T cell donors, *PRDX1*-knockout T cells were generated from three additional healthy donors and cultured in either ascites or media. In contrast to the screening T cell donor (donor 1), *PRDX1*-knockout did not consistently enhance IFN-γ production across additional healthy donors in ascites and media (donor 3, 4, 5) (Fig. S3F-G), highlighting inter-donor heterogeneity in the functional consequences of *PRDX1*-knockout. Consistent with the validation experiments using the pooled screen donor (Fig. 3 and 4), mitochondrial membrane potential remained largely unchanged following *PRDX1*-knockout in both ascites and media conditions in the three additional donors tested (Fig. S3H-I).

As PRDX1 expression may also contribute to donor-specific phenotypes, relative PRDX1 mRNA expression was quantified in stimulated T cells from donors 1 (CRISPR screening donor, Fig. 3 and 4), 3, 4, and 5 cultured in HGSOC ascites supernatant. PRDX1 transcript abundance varied slightly between donors, with donor 5 exhibiting the highest PRDX1 expression (Fig. S3J). Together, these findings suggest endogenous PRDX1 expression remains consistent between T cell donors, apart from donor 5, and is therefore unlikely to explain the biological heterogeneity in *PRDX1* knockout responses.

Given the inter-patient heterogeneity of the HGSOC tumor microenvironment, the effects of *PRDX1*-knockout were next examined in T cells cultured in ascites supernatants from multiple patients. The impact of *PRDX1*-knockout on IFN-γ production varied between ascites donors, with more modest effects observed in the additional samples (Fig. S3K) compared to the sample used in the pooled CRISPR screen (Fig. 4C). In line with previous results, *PRDX1*-knockout did not consistently alter mitochondrial membrane potential across any of the additional ascites conditions examined (Fig. S3L).

### PRDX1 deletion enhances T cell activation and metabolic fitness

To further characterize the functional consequences of *PRDX1* deletion, we next examined T cell, proliferation, activation, inflammatory cytokine production, viability and metabolic phenotypes following T cell receptor (TCR) restimulation in either complete media or HGSOC ascites (Fig. S4A). Although *PRDX1*-knockout cells displayed comparable expansion to control cells throughout most of the culture period, a modest reduction in cumulative expansion was observed by day 12 (Fig. 5A).

**Fig. 5.**
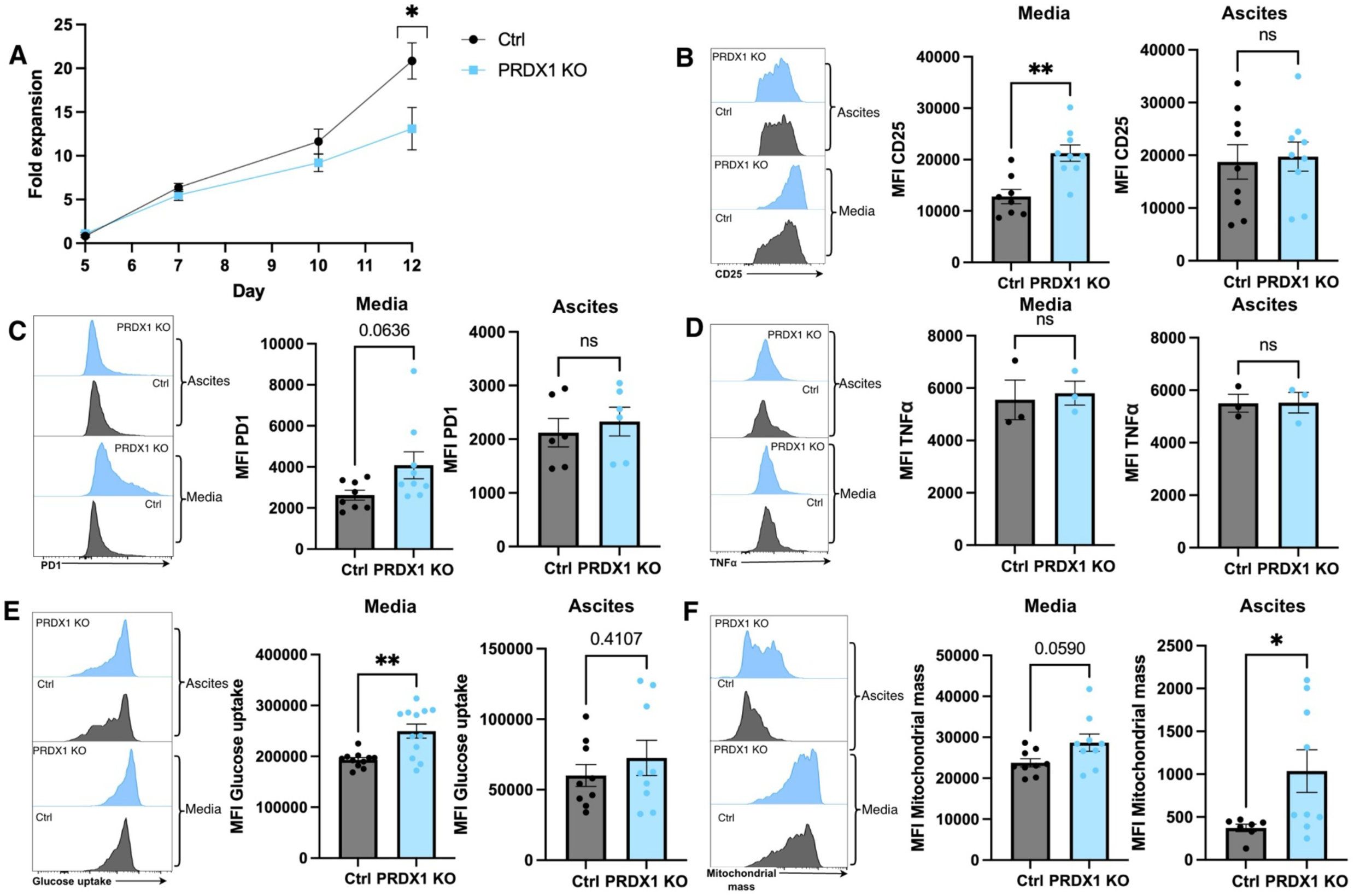
*PRDX1* deletion enhances T cell activation and metabolic phenotypes. **(A)** Expansion of control and *PRDX1*-knockout T cells during in vitro culture. Fold expansion was measured from day 5 to day 12 after activation. **(B)** CD25 expression following *PRDX1* deletion. Representative histograms and quantification of CD25 mean fluorescence intensity (MFI) are shown for cells stimulated for 48 hrs in either media or HGSOC ascites. **(C)** PD-1 expression following *PRDX1* deletion. Representative histograms and quantification of PD-1 MFI are shown for cells stimulated for 48 hrs in media or HGSOC ascites. **(D)** TNFα production following *PRDX1* deletion. Representative histograms and quantification of intracellular TNFα are shown after 48 hr stimulation in media or HGSOC ascites. **(E)** Glucose uptake following *PRDX1* deletion. Glucose uptake was measured after 48 hr stimulation in media or HGSOC ascites. **(F)** Mitochondrial mass following PRDX1 deletion. Representative histograms and quantification of mitochondrial mass are shown after 48 hr stimulation in media or HGSOC ascites. For panels (A–F), T cells were cultured in media or ascites derived from the same HGSOC patient sample used in the pooled CRISPR screen (Fig. 1I-J: patient 3). Control T cells represent cells treated with no sgRNA, intergenic sgRNA, or *AAVS1*-targeting sgRNA; *PRDX1*-knockout T cells represent cells treated with sgRNA 1, sgRNA 8, and sgRNA 10 (from Fig. 3B). Each point represents a distinct sgRNA tested in a single T cell donor; some sgRNAs were evaluated in technical replicates across multiple experiments. Bars represent mean ± SEM. Statistical significance was determined using Welch’s t-test. *P < 0.05, **P < 0.01; ns, not significant.

Consistent with the enhanced IFN-γ phenotype observed previously, *PRDX1*-knockout T cells exhibited increased expression of the activation marker CD25 under media conditions, whereas CD25 expression was unchanged following culture in ascites (Fig. 5B). Similarly, PD-1 expression showed a trend toward increased expression in media but was not altered in ascites 48 hours after restimulation (Fig. 5C). In contrast, intracellular TNFα production was unaffected by *PRDX1* deletion under either condition (Fig. 5D), indicating that *PRDX1* deletion selectively influences aspects of T cell activation rather than broadly augmenting inflammatory cytokine production.

As alterations in cellular metabolism can influence T cell survival, the effect of *PRDX1* deletion on T cell viability was assessed next. *PRDX1*-knockout had no effect on viability in complete media but increased the viability of T cells cultured in HGSOC ascites (Fig. S4B), suggesting that loss of PRDX1 confers a survival advantage under suppressive ascites conditions.

Given the metabolic role of PRDX1, the effect of PRDX1 deletion on glucose uptake and mitochondrial mass was examined. *PRDX1*-knockout increased glucose uptake under media conditions and exhibited a similar trend in ascites (Fig. 5E). Conversely, mitochondrial mass was increased in ascites-cultured T cells and trended toward an increase in media following *PRDX1* deletion (Fig. 5F). Together, these findings demonstrate that *PRDX1* deletion promotes metabolic remodeling characterized by enhanced glucose uptake and increased mitochondrial quantity. To determine whether these metabolic phenotypes differed between T cell subsets, CD4⁺ and CD8⁺ T cells were analyzed separately (Fig. S4C-F). Under ascites and media conditions, the increase in glucose uptake upon *PRDX1* deletion was primarily restricted to CD8⁺ T cells, whereas CD4⁺ T cells were unaffected (Fig. S4C-D). In contrast, *PRDX1* deletion increased mitochondrial mass in both CD4⁺ and CD8⁺ T cells under ascites conditions (Fig. S4E), whereas mitochondrial mass remained unchanged under media conditions (Fig. S4F). These subset-specific analyses suggest that enhanced glucose uptake is predominantly a feature of CD8⁺ T cells, whereas increased mitochondrial content occurs across both major T cell populations.

### PRDX1 deletion differentially regulates CAR-T cell cytotoxicity across donors

To determine whether the functional effects of *PRDX1* deletion extended to chimeric antigen receptor (CAR) T cells, primary human T cells were transduced with a folate receptor α (FRα)-specific CAR (*36*) followed by sequential CRISPR-Cas9-mediated *PRDX1* deletion and intracellular staining for IFN-γ. Consistent with previous observations, *PRDX1*-knockout T cells exhibited an increased frequency of IFN-γ-producing cells following TCR-mediated restimulation in media (Fig. 6A). However, this enhancement was not observed following stimulation of CAR-T cells, either through the TCR or CAR, where *PRDX1* deletion had no significant effect on IFN-γ production (Fig. 6A).

**Fig. 6.**
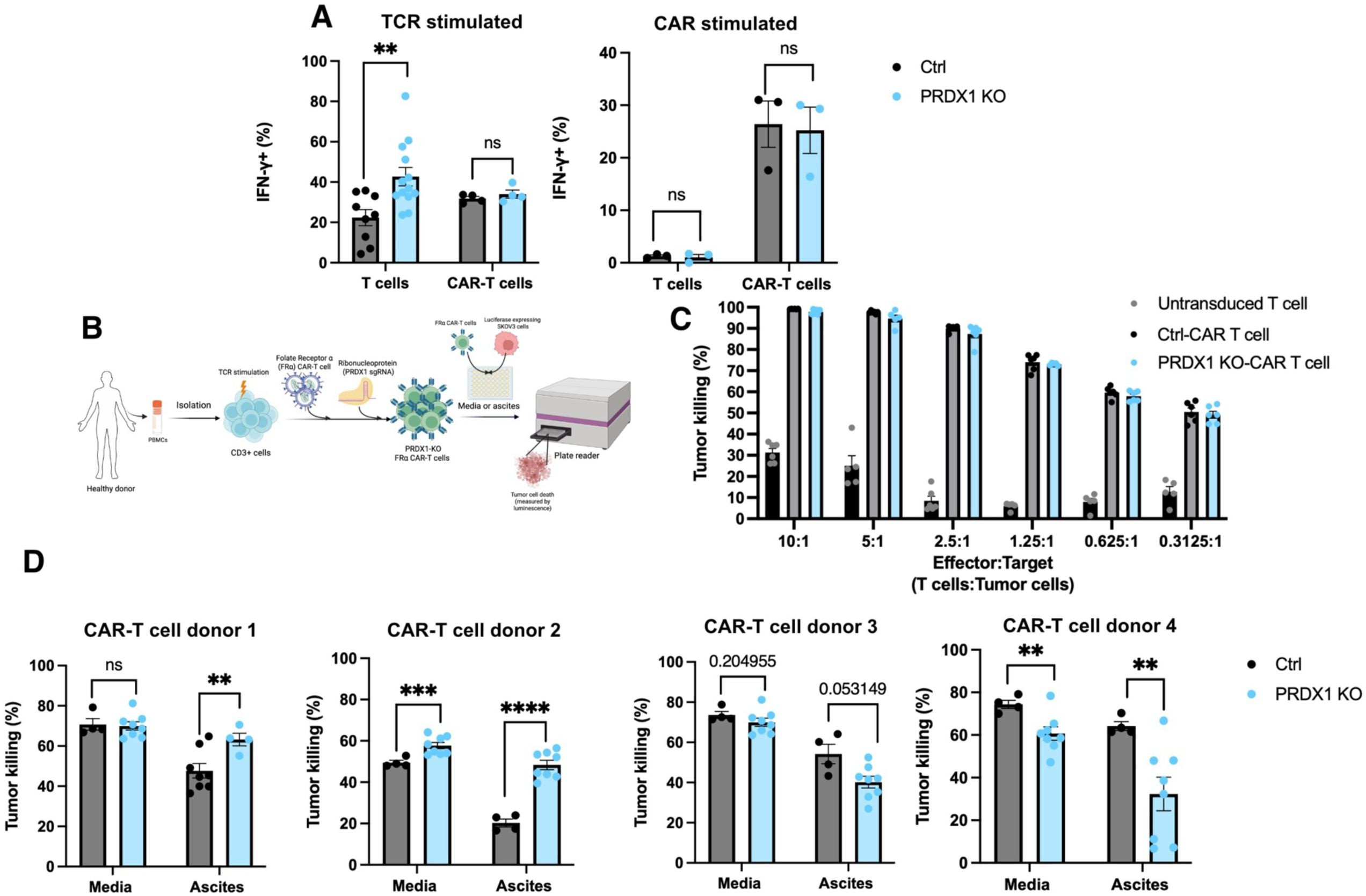
*PRDX1* deletion differentially affects CAR-T cell cytotoxicity across donors. **(A)** IFN-γ production by control and *PRDX1*-knockout untransduced T cells and FRα CAR-T cells following either TCR-or CAR-mediated stimulation. Cells were co-cultured with FRα-expressing SKOV3 target cells (CAR stimulated), and IFN-γ production was assessed by intracellular cytokine staining. **(B)** Schematic of the experimental workflow used to generate *PRDX1*-knockout FRα CAR-T cells and evaluate cytotoxic function. CD3+ T cells were isolated from healthy donors, activated, transduced with an FRα-targeting CAR on day 1, enriched by CAR-positive bead sorting on day 4, and electroporated with *PRDX1*-targeting ribonucleoprotein complexes on day 6. Edited CAR-T cells were cultured in media or HGSOC ascites (patient 1, Fig. 1I-J) and co-cultured with luciferase-expressing SKOV3 target cells. Tumor cell killing was quantified by loss of luminescence. Created with BioRender. **(C)** Cytotoxic activity of control and *PRDX1*-knockout FRα CAR-T cells against SKOV3-luciferase target cells across a range of effector-to-target (E:T) ratios under standard media conditions. Data are shown for a single T cell donor. **(D)** Cytotoxic activity of control and *PRDX1*-knockout FRα CAR-T cells following a 24 hr co-culture with SKOV3-luciferase target cells at an E:T ratio of 1:1 in either media or HGSOC ascites. Data are shown for four independent CAR-T cell donors. For figures A-B and C-D, control T cells represent cells treated with no sgRNA, intergenic sgRNA, or *AAVS1*-targeting sgRNA; *PRDX1*-knockout T cells represent cells treated with sgRNA 1, sgRNA 8, and sgRNA 10 (from Fig. 3B). Each point represents a distinct sgRNA tested in one donor sample; some sgRNAs were evaluated in technical replicates across multiple experiments. Bars represent mean ± SEM. Statistical significance was determined using Welch’s t-test. **P < 0.01, ***P < 0.001, ****P < 0.0001; ns, not significant.

The effect of *PRDX1* deletion on CAR-T cell cytotoxicity against FRα-expressing SKOV3 tumor cells was next assessed (Fig. 6B). Across a range of effector-to-target ratios under media conditions, both control and *PRDX1*-knockout CAR-T cells exhibited comparable tumor-killing capacity, with no consistent differences observed between groups (Fig. 6C).

Because previous experiments demonstrated that PRDX1 phenotypes were influenced by T cell donors, CAR-T cell cytotoxicity was next evaluated in multiple healthy donors in media and suppressive HGSOC ascites supernatant (patient 1 ascites, Fig. 1J). Under media conditions, *PRDX1* deletion produced variable effects across T cell donors, enhancing cytotoxicity in donor 2, reducing cytotoxicity in donor 4, and having minimal effect in donors 1 and 3 (Fig. 6D). Greater donor-dependent heterogeneity emerged under ascites conditions. *PRDX1* deletion significantly enhanced tumor killing in donors 1 and 2, showed a trend toward reduced cytotoxicity in donor 3, and markedly impaired cytotoxicity in donor 4 (Fig. 6D). Collectively, these findings indicate that the effects of *PRDX1* deletion on CAR-T cell cytotoxicity are highly context- and donor-dependent.

### PRDX1 deletion increases intracellular ROS and impairs autophagic flux in primary human T cells

To investigate the mechanisms underlying the phenotypic effects of *PRDX1* deletion, intracellular reactive oxygen species (ROS) were first assessed following TCR restimulation. As expected, based on the established antioxidant function of PRDX1 (*34*, *35*), *PRDX1*-knockout T cells exhibited significantly increased intracellular ROS compared with control cells (Fig. 7A), confirming that PRDX1 disruption alters cellular redox homeostasis.

**Fig. 7.**
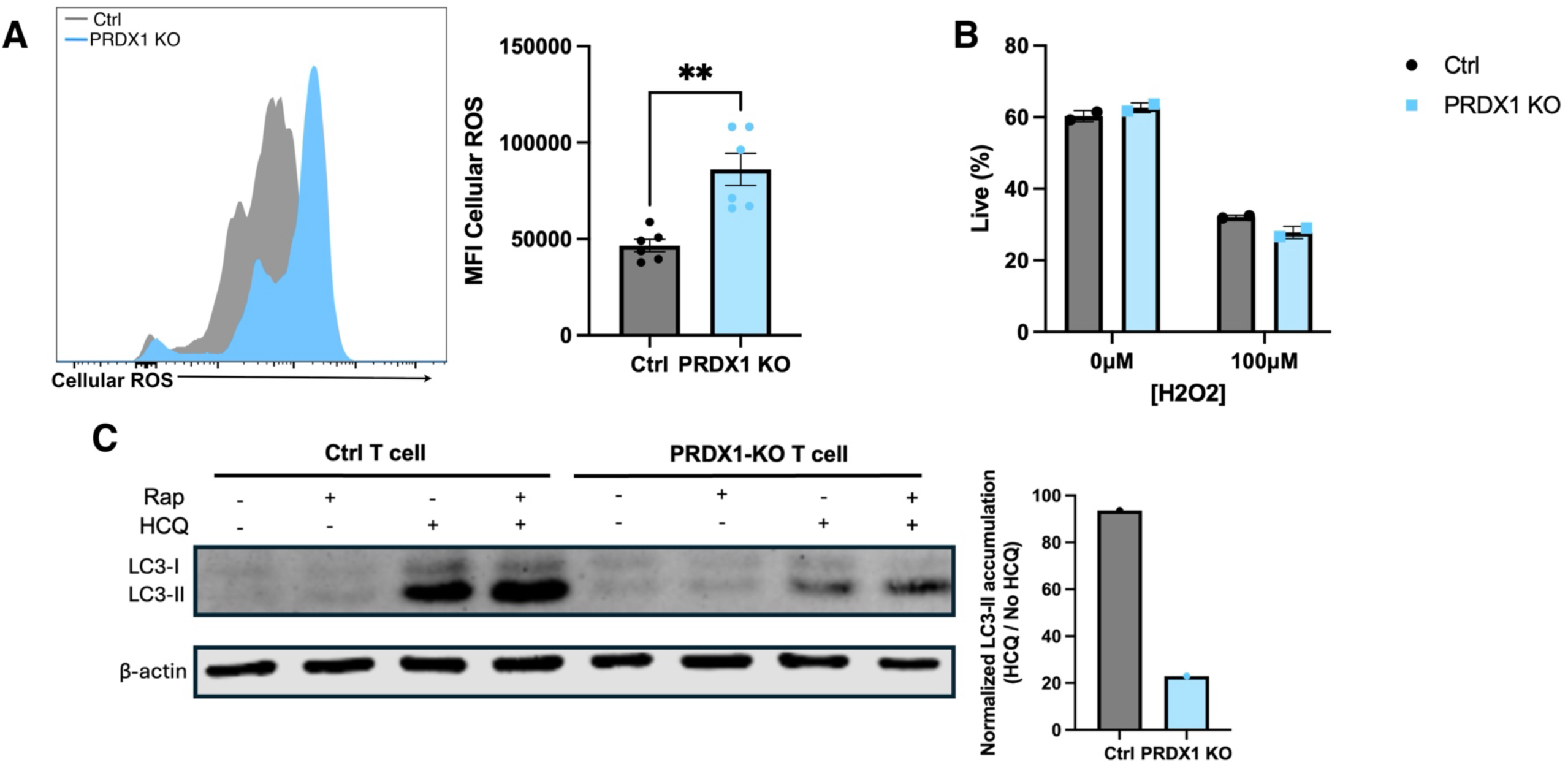
*PRDX1* deletion increases cellular ROS and reduces autophagic flux in primary human T cells. **(A)** Cellular reactive oxygen species (ROS) following *PRDX1* knockout. Cellular ROS was measured after 48 hr stimulation in media. Each point represents a distinct sgRNA tested in one donor sample. Control samples included no sgRNA, intergenic sgRNA, or *AAVS1*-targeting sgRNA, whereas PRDX1-knockout samples included PRDX1 sgRNA 1, sgRNA 8, or sgRNA 10 (from Fig. 3B). Some sgRNAs were evaluated in technical replicates across multiple experiments. Statistical significance was determined using Welch’s t-test. **(B)** Sensitivity of control and PRDX1-knockout T cells to oxidative stress. Cells were cultured in the presence or absence of 100 μM H₂O₂ for 48 hrs, and viability was assessed. Data are shown for one T cell donor, with each point representing a technical replicate. Control samples consisted of a pooled mixture of cells transfected with no sgRNA, an *AAVS1*-targeting sgRNA, or an intergenic sgRNA, whereas *PRDX1*-knockout samples consisted of a pooled mixture of cells transfected with PRDX1-targeting sgRNAs 1, 8, and 10 (from Fig. 3B). **(C)** Assessment of autophagic flux following *PRDX1* knockout. Representative immunoblot showing LC3-I and LC3-II levels in control and *PRDX1*-knockout T cells following treatment with rapamycin (Rap; 400 nM), hydroxychloroquine (HCQ; 30 μM), or both for 20 hrs. LC3-II accumulation was quantified following normalization to β-actin and expressed as the ratio of HCQ-treated to untreated samples. For A-B, bars represent mean ± SEM. Statistical significance was determined using Welch’s t-test. **P < 0.01.

To determine whether elevated ROS rendered *PRDX1*-knockout T cells more susceptible to oxidative stress, cells were treated with 100μM H₂O₂. Both control and *PRDX1*-knockout T cells exhibited a comparable reduction in viability, indicating that *PRDX1* deletion did not measurably alter sensitivity to exogenous oxidative stress under these conditions (Fig. 7B).

To determine whether the phenotype observed following PRDX1 deletion was unique to PRDX1 or shared among other members of the peroxiredoxin family, *PRDX2*, *PRDX3*, and *PRDX4* were individually deleted in primary human T cells. TIDE analysis confirmed efficient gene editing for all three targets, with indel frequencies ranging from approximately 50–80% (Fig. S5A). Intracellular ROS levels were determined following 48 hours of T cell stimulation in media to determine whether deletion of these family members altered ROS accumulation. Similar to *PRDX1*, deletion of *PRDX2* significantly increased cellular ROS levels compared with control T cells (Fig. S5B). *PRDX4* deletion also significantly elevated cellular ROS, whereas *PRDX3* deletion did not significantly alter ROS levels but exhibited a trending increase (Fig. S5B). We next examined whether these changes in redox homeostasis translated into enhanced T cell effector function. In HGSOC ascites, only *PRDX2* deletion significantly increased the frequency of IFN-γ-producing T cells compared with controls (Fig. S5C). In contrast, deletion of *PRDX3* or *PRDX4* did not significantly alter IFN-γ production. Likewise, under media conditions, deletion of *PRDX2*, *PRDX3*, or *PRDX4* had no significant effect on the proportion of IFN-γ-producing T cells (Fig. S5D). Together, these findings demonstrate that PRDX family members differentially regulate T cell redox homeostasis and effector function. Although deletion of both *PRDX2* and *PRDX4* increased intracellular ROS, only *PRDX2* deletion enhanced IFN-γ production under suppressive ascites conditions, suggesting that elevated ROS alone is insufficient to promote improved T cell effector function.

Because PRDX1 and ROS have been implicated in regulating autophagy (*33*, *37*), autophagic flux was next examined by monitoring LC3-II accumulation following inhibition of lysosomal degradation with hydroxychloroquine (HCQ). As expected, HCQ treatment increased LC3-II accumulation in control T cells, whereas rapamycin-induced autophagy further enhanced LC3-II levels (Fig. 7C). In contrast, *PRDX1*-knockout T cells exhibited markedly reduced LC3-II accumulation following HCQ treatment, despite comparable β-actin loading (Fig. 7C). Quantification demonstrated substantially reduced HCQ-induced LC3-II accumulation in *PRDX1*-knockout cells relative to controls, consistent with impaired autophagic flux (Fig. 7C).

Collectively, these findings demonstrate that *PRDX1* deletion increases intracellular ROS while impairing autophagic flux, identifying dysregulated redox homeostasis and autophagy as potential mechanisms underlying the functional phenotypes observed following *PRDX1* deletion.

## Discussion

Durable responses to T cell-based immunotherapy for HGSOC remain limited. A key contributor is the metabolically restricted tumor microenvironment (TME), which has been shown to have low levels of essential nutrients and high abundance of immunosuppressive metabolites (*9*, *13*, *26*, *38*). Numerous CRISPR-based genetic screens have identified regulators of T cell function, yet the majority have relied on murine models or simplified in vitro systems that do not physiologically recapitulate the complexity of the human TME (*15*, *17*, *28*, *39*, *40*). Here, we developed a pooled CRISPR-Cas9 screening platform performed in primary human T cells cultured in patient HGSOC ascites, enabling the identification of metabolic regulators that influence T cell function within a relevant suppressive TME. While the ascites environment differs from the primary tissue in HGSOC, isolation of sufficient quantities of tumor-interstitial fluid is a significant practical challenge. Moreover, several groups have shown that infusion of both systemic- and cell-based infusion into the ascites is a viable, safe, and effective site of delivery (*41*, *42*). Using this CRISPR-based screening strategy, peroxiredoxin 1 (*PRDX1)* was discovered as a negative regulator of T cell effector function. We demonstrate that loss of PRDX1 enhances the frequency of IFN-γ-producing T cells without substantially affecting mitochondrial membrane potential. Importantly, the magnitude of the *PRDX1*-knockout phenotype was influenced by both T cell donor and ascites donor, highlighting the context-dependent nature of the phenotype uncovered in the pooled CRISPR screen.

Human ascites contains a complex mixture of soluble cytokines, metabolites, lipids, extracellular vesicles, and other immunomodulatory factors that collectively shape T cell function (*22*, *23*, *43*). Although reductionist culture systems have proven valuable for dissecting individual pathways, they cannot fully reproduce the multifactorial nature of immune suppression encountered within human tumors (*40*). The use of primary patient ascites therefore represents an important strength of this screening platform, as it exposes T cells to clinically relevant, patient-specific features of the HGSOC microenvironment that are not faithfully reproduced in conventional culture systems or murine models. Performing the screen within this complex human-derived environment may improve the translational relevance of the identified regulators while also capturing the substantial interpatient heterogeneity characteristic of HGSOC. However, ascites does not fully recapitulate the complexity of an intact solid tumor. Unlike tumor tissue, ascites lacks structural and mechanical barriers and is generally nutrient replete, and therefore primarily models the soluble and metabolic components of the HGSOC microenvironment. Accordingly, complementary in vivo studies will be required to determine whether the regulators identified using this platform similarly influence T cell function within intact solid tumors and whether these effects extend to other malignancies.

An important observation throughout this study was the substantial variability observed between healthy donors and ascites samples. Rather than representing technical variability, we believe this heterogeneity reflects an important feature of human biology. Human T cells differ considerably between individuals because of genetic background, environmental exposures, antigen experience, and differentiation state (*44*, *45*). Likewise, ovarian cancer ascites exhibits marked patient-to-patient variability in cytokine composition, metabolic profile, and immunosuppressive capacity (*43*). Consequently, it is perhaps unsurprising that PRDX1 perturbations did not produce identical phenotypes across every donor or every ascites sample. Instead, these findings suggest that the consequences of manipulating metabolic pathways are highly dependent on both the intrinsic properties of the responding T cell and the surrounding microenvironment. Furthermore, this metabolic heterogeneity is consistent with the non-uniform distribution of genomic alterations and neoantigens across tumors and represents an additional layer of tumor heterogeneity beyond the well-established genomic diversity reported in the literature (*46*, *47*).

Among the candidates identified in the screen, *PRDX1* emerged as one of the highest-confidence hits, owing to its strong statistical significance and the consistent enrichment of multiple independent sgRNAs within the highly functional T cell population. Validation studies confirmed efficient gene disruption and PRDX1 protein depletion and demonstrated that *PRDX1* deletion increased the frequency of interferon gamma (IFN-γ)-producing T cells, supporting its role as a negative regulator of T cell effector function. Beyond this phenotype, *PRDX1*-knockout promoted a broader metabolic and activation program, including increased glucose uptake, elevated mitochondrial mass, enhanced CD25 expression, and improved viability under ascites conditions, while having minimal effects on tumor necrosis factor alpha (TNFα) production or mitochondrial membrane potential. Collectively, these findings suggest that PRDX1 influences multiple aspects of T cell activation and metabolic adaptation.

The increase in glucose uptake following *PRDX1* deletion may be particularly relevant in the metabolically restrictive TME, where T cells must compete with tumor cells for limited nutrients (*48*, *49*). Enhanced glucose uptake could provide *PRDX1*-knockout T cells with a competitive advantage by improving access to a key substrate required for activation, cytokine production, and effector function (*50*). In parallel, the increase in mitochondrial mass observed after *PRDX1* deletion may reflect an altered mitochondrial fitness state. This is consistent with studies linking mitochondrial biogenesis and mitochondrial mass to improved T cell and CAR-T cell function (*51*, *52*). In CLL, malignant cells impair CD8⁺ T cell mitochondrial fitness and limit CAR-T efficacy, supporting the idea that mitochondrial dysfunction can constrain cellular immunotherapy responses (*52*). Conversely, higher mitochondrial mass in pre-lymphodepletion T cells has been associated with early response to CAR-T therapy in relapsed/refractory non-Hodgkin lymphoma (*51*). Together, these findings suggest that the metabolic changes observed following *PRDX1* deletion, including increased glucose uptake and mitochondrial mass, may contribute to improved T cell fitness in select suppressive microenvironments.

Although *PRDX1* deletion consistently increased mitochondrial mass, it did not significantly alter mitochondrial membrane potential (ΔΨm). While these measurements are often interpreted together, they reflect distinct aspects of mitochondrial biology. Mitochondrial mass quantifies total mitochondrial content, whereas ΔΨm reflects the electrochemical potential across the inner mitochondrial membrane and serves as an indicator of mitochondrial bioenergetic activity on a per-mitochondrion basis (*53*, *54*). Consequently, an increase in mitochondrial content does not necessarily correspond to increased membrane potential if individual mitochondria maintain similar bioenergetic function. One potential explanation for our findings is that impaired autophagic flux following *PRDX1* deletion reduces mitochondrial turnover, resulting in the accumulation of mitochondria without substantially altering their membrane potential (*55*). This interpretation is consistent with the observed increase in mitochondrial mass together with preserved ΔΨm and supports the notion that mitochondrial quantity and mitochondrial function can be independently regulated.

Interestingly, enhanced cytokine production did not uniformly translate into improved CAR-T cell function. While *PRDX1* deletion consistently enhanced IFN-γ production following conventional TCR stimulation in the screening T cell donor, CAR-mediated activation was largely unaffected, and cytotoxic activity varied substantially between donors under ascites conditions. These findings emphasize that cytokine production and tumor killing represent distinct functional outputs governed by overlapping but non-identical regulatory pathways. Furthermore, these results raise the possibility that CAR signaling architecture modifies downstream signaling networks such that the functional effects of *PRDX1* deletion are attenuated. Therefore, genetic modifications that improve one aspect of T cell function may not universally enhance antitumor activity, particularly when combined with additional genetic engineering approaches such as CAR expression.

Mechanistically, *PRDX1* deletion increased intracellular reactive oxygen species (ROS), consistent with the established role of PRDX1 as a key antioxidant enzyme. Elevated ROS has complex effects on T cell biology. Moderate increases in ROS can promote T cell receptor (TCR) signaling and effector differentiation, whereas excessive ROS contributes to oxidative damage, metabolic dysfunction, and impaired persistence (*56*). Therefore, *PRDX1* deletion may only be beneficial in environments where there is relatively low oxidative burden. Given that deletion of other PRDX family members similarly increased ROS production but did not consistently enhance IFN-γ production, we reasoned that elevated ROS alone is insufficient to explain the increased effector function observed following *PRDX1* knockout.

While autophagy has traditionally been viewed as essential for maintaining T cell homeostasis and long-term survival, emerging evidence indicates that transient repression of autophagic flux accompanies antigen-driven activation and can facilitate effector differentiation (*57–59*). Recent work demonstrated that TCR signaling and inflammatory cytokines rapidly suppress autophagy in activated CD8⁺ T cells, thereby preserving cytolytic effector molecules, nutrient transporters, and metabolic machinery required for effective immune responses (*59*). Likewise, genetic inhibition of autophagy has been shown to enhance antitumor activity of CD8⁺ T cells in certain settings, suggesting that reduced autophagic flux is not universally detrimental but may instead represent a physiological feature of highly activated effector T cells (*60*). Accordingly, the impaired autophagic flux observed following *PRDX1* deletion may contribute to the enhanced effector phenotype observed in our screen and validation experiments. Although the precise relationship between redox regulation, autophagy, and T cell effector function remains to be determined, our findings identify these pathways as plausible downstream mechanisms through which PRDX1 regulates T cell biology.

Notably, the pooled CRISPR screen was performed using T cells from a single healthy donor cultured in ascites from a single HGSOC patient. Although subsequent validation experiments incorporated multiple healthy donors and ascites samples, performing arrayed screens to validate pooled screen hits across numerous donor– patient combinations remains technically prohibitive. Second, while mechanistic studies demonstrated that *PRDX1* deletion increased intracellular ROS and impaired autophagic flux, these experiments do not establish a causal relationship between either pathway and the enhanced effector function observed following *PRDX1* deletion in T cells. Future studies will be required to determine whether modulation of ROS, autophagy, or other downstream pathways directly mediates the functional phenotype.

Here, we establish a pooled CRISPR screening strategy that combines primary human T cells with a patient-derived physiological model of the ovarian cancer TME. This approach captures biologically meaningful heterogeneity that is absent from many existing screening platforms and allows for the identification of regulators of human T cell function under clinically relevant suppressive conditions. Beyond identifying PRDX1 as a context-dependent regulator of T cell function, our findings highlight the importance of incorporating human donor and patient variability into functional genomic studies to improve the translational relevance of discoveries aimed at enhancing cancer immunotherapy.

## Materials and Methods

### Human peripheral blood samples

Peripheral blood mononuclear cells (PBMCs) were isolated from healthy donor leukopaks (STEMCELL Technologies) by density-gradient centrifugation using Ficoll-Paque. CD3+ T cells were isolated from PBMCs using Miltenyi MACS separation protocol, according to the manufacturer’s instructions. Isolated T cells were used for pooled CRISPR screening, validation experiments, metabolic phenotyping, CAR-T cell generation and autophagy flux assays, as described below. Each T cell donor was used independently, and biological replicates represent experiments performed using cells isolated from different donors. Human samples were permitted for use under Research Ethics Board (REB) protocols approved by the University of Victoria (human ethics: 19-0067) and the University of British Columbia (Biosafety: B23-0067; human ethics H25-00625).

### Patient ascites samples

Ascites samples were collected from patients with high-grade serous carcinoma (HGSOC) undergoing clinical management at BC Cancer (Victoria). Ascites samples were processed by centrifugation to remove cellular material, and the clarified supernatant was collected and stored at -80°C until use. All ascites samples were collected prior to any treatment of patients. For flow cytometry experiments, prior to T cell culture, ascites supernatant was thawed and used without further modification. For CAR-T cell cytotoxicity assays, ascites supernatant was thawed and filtered using a 40µm cell strainer. Biospecimens were obtained with assistance from the IROC-Tumor Tissue Repository Biobank (BC Cancer, Victoria). Patient samples were permitted for use under Research Ethics Board (REB) protocols approved by the University of Victoria (human ethics: 19-0067) and the University of British Columbia (Biosafety: B23-0067; human ethics H25-00625).

Patient-derived ascites was used as an *ex vivo* model of the ovarian tumor microenvironment throughout this study. All experiments using ascites were performed using 100% ascites supernatant. The ascites sample from patient 3 (Fig. 1 I-J) was used for pooled CRISPR-Cas9 screening and the majority of validation experiments, while additional ascites samples from independent HGSOC patients were used to assess the reproducibility of observed phenotypes. A separate highly suppressive ascites sample from patient 1 (Fig. 1 I-J) was used for CAR-T cell cytotoxicity experiments.

Patient information is included in Table S1.

### Untargeted metabolomic profiling

Untargeted metabolomic profiling was performed on patient-matched serum and ascites supernatant samples obtained from three individuals with HGSOC. Serum and ascites samples were collected prior to any treatment of patients. Serum was processed using serum separator tubes, aliquoted, and stored at −80°C prior to sending for metabolite analysis. Ascites samples were processed by centrifugation to remove cellular material, and the clarified supernatant was collected and stored at -80°C until sending for metabolite analysis. Metabolites were extracted and analyzed by liquid chromatography-mass spectrometry (LC-MS) by Agios to identify over 3000 metabolites.

Raw data were processed using MetaboAnalyst. Peak intensity data were uploaded and analyzed using the Statistical Analysis (one-factor) module for heatmap generation or the Metadata Table module for principal component analysis (PCA). Data were filtered using the interquartile range (IQR) method, normalized by sum, log-transformed, and auto-scaled prior to downstream analyses. Heatmaps were generated using Euclidean distance with average linkage clustering for one-factor analyses.

### Primary human T cell activation and culture

Following isolation, T cells were cultured in complete ImmunoCult™-XF T Cell Expansion Medium (STEMCELL; 10981) supplemented with 1% penicillin-streptomycin and 300U/mL recombinant human IL-2 (PeproTech; 20002100UG). Cells were expanded according to the manufacturer’s protocol for ImmunoCult™-XF T Cell Expansion Medium (STEMCELL Technologies). Cells were maintained at 37°C in a humidified incubator with 5% CO₂.

For activation following isolation from PBMCs (day 0), T cells were activated using ImmunoCult anti-CD3/CD28 activation reagent (STEMCELL Technologies; 10971) at 25μL/mL. For reactivation following expansion (day 12 for 48 hr stimulation; day 11 for 10 hr stimulation), T cells were activated using ImmunoCult anti-CD3/CD28/CD2 activation reagent (STEMCELL Technologies; 10970) at 25μL/mL. Unless otherwise indicated, cells were expanded in media for 12 days prior to downstream assays. Unless otherwise indicated, for experiments evaluating T cell function and metabolism, on day 12 post-isolation, T cells were transferred into either complete media or 100% patient HGSOC ascites supernatant and reactivated with anti-CD3/CD28/CD2 for 48 hrs before phenotypic analysis.

### Flow cytometry

Flow cytometry was used to assess T cell activation, metabolism, and effector function. Following incubation, cells were washed with phosphate-buffered saline (PBS) and stained according to standard protocols.

Metabolic phenotyping was performed using fluorescent probes specific for mitochondrial membrane potential (MitoTracker Deep Red, Thermo Fisher, M46753), mitochondrial mass (MitoTracker Green, Thermo, M46750), glucose uptake (GlucoseCy5, Sigma, SML3233-5MG), and cellular reactive oxygen species (Total ROS Detection Reagent, Thermo Fisher, 88-5930-74). Cells were incubated with each probe according to the manufacturer’s recommendations before washing and either continuing with staining protocol (viability and surface marker staining) or immediate flow cytometric analysis.

Cell viability was assessed using eFluor™ 506 Fixable Viability Dye (Thermo Fisher, 65-0866-18) according to the manufacturer’s instructions. Subsequently, for surface marker analysis, cells were stained with fluorophore-conjugated antibodies against CD4 (BioLegend, 300526), CD8 (BioLegend, 301030), CD25 (Thermo Fisher, 46-0257-41) and PD-1 (BioLegend, 329950) for 30 minutes at 4°C in the dark. Following staining, cells were washed with PBS supplemented with 3% human serum and analyzed immediately.

For intracellular cytokine analysis, cells were fixed and permeabilized using the BD Cytofix/Cytoperm Fixation/Permeabilization Kit, according to the manufacturer’s instructions, before staining with fluorophore-conjugated antibodies against interferon-γ (IFN-γ) (BD, 554552) and tumor necrosis factor alpha (TNFα) (BD, 554512) (and against surface markers, if applicable). Intracellular cytokine production was quantified following 48 hrs or 10 hrs of T cell receptor (TCR) restimulation.

Flow cytometry data were acquired on a Cytek Aurora spectral flow cytometer and analyzed using FlowJo version 10.8.2. Lymphocytes, representing an enriched population of isolated CD3⁺ T cells, were identified based on forward- and side-scatter characteristics, followed by exclusion of doublets and non-viable cells where appropriate. Subsequent gating strategies were tailored to each panel. For pooled CRISPR screening experiments, after lymphocyte, singlet, and viability gating, cells were additionally gated on the highest and lowest 20% mitochondrial membrane potential and IFN-γ expression to isolate high-effector function (IFN-γ-high; ΔΨm-high) and low-effector function (IFN-γ-low; ΔΨm-low) populations for fluorescence-activated cell sorting (FACS).

All flow cytometry antibodies can be found in Table S2.

### Lentivirus production

Lentivirus was produced in HEK293T cells using a third-generation packaging system and Lipofectamine 3000 (Thermo Fisher, L3000015). HEK293T cells were expanded in Dulbecco’s Modified Eagle Medium (DMEM, Cytiva, SH3002201) supplemented with 10% fetal bovine serum and 1% penicillin-streptomycin until sufficient cells were obtained for virus production. For each lentiviral construct, 3.3×10^7^ HEK293T cells were seeded into T175 flasks in Opti-MEM I Reduced Serum Medium supplemented with GlutaMAX (Gibco, 51985034) and 5% FBS one day prior to transfection to achieve approximately 95–100% confluency at the time of transfection.

For transfection, transfer plasmid DNA was co-transfected with the packaging plasmid psPAX2 (Addgene #12260) and the envelope plasmid pMD2.G (Addgene #12259) using Lipofectamine 3000 (Thermo Fisher, L3000015) according to the manufacturer’s protocol. For each T175 flask, 14 μg transfer plasmid, 28 μg psPAX2, and 14 μg pMD2.G were diluted in Opti-MEM together with P3000 reagent (Thermo Fisher, L3000015), while Lipofectamine 3000 (Thermo Fisher, L3000015) was diluted separately in Opti-MEM (Gibco, 31985062). Following a 15 min incubation at room temperature to allow DNA-lipid complex formation, the complexes were added dropwise to HEK293T cells. Six hours after transfection, fresh complete Opti-MEM medium was added to restore the culture volume to 37mL.

Virus-containing supernatants were collected at 24 and 48 hrs post-transfection. Following the 24 hr harvest, fresh complete Opti-MEM medium was added to the cultures before the final harvest at 48 hr. Cell debris was removed by centrifugation at 2,000 rpm for 10 min, followed by filtration through a 0.45 μm syringe filter. The 24 hr and 48 hr viral harvests for each construct were combined prior to concentration. Lentiviral particles were concentrated by ultracentrifugation at 25,000 rpm (approximately 90,000 × g) for 90 min at 4°C using an Optima XL-100K ultracentrifuge equipped with an SW32 Ti rotor (Beckman Coulter). Following centrifugation, the supernatant was carefully removed and viral pellets were resuspended in Opti-MEM. Virus from the same construct was pooled, aliquoted into sterile microcentrifuge tubes, and stored at −80°C until use.

### Lentiviral titre determination

The functional titre of the pooled CRISPR sgRNA lentiviral library was determined in primary human T cells to identify the volume of virus required to achieve a multiplicity of infection (MOI) of 0.3. Primary human CD3⁺ T cells were isolated from healthy donor PBMCs by magnetic separation (Miltenyi Biotec, 130-050-101), activated with CD3/CD28 activators (STEMCELL Technologies, 10971) in ImmunoCult-XF medium (STEMCELL Technologies, 10981) supplemented with 300 U/mL IL-2 (PeproTech, 200-02-100UG), and seeded at 1×10^5^ cells per well in 96-well round-bottom plates.

A 3.16-fold serial dilution of concentrated lentiviral supernatant was prepared in Opti-MEM (Gibco, 31985062) and added to activated T cells. Twenty-four hours after transduction, cells were cultured in the presence or absence of 2 μg/mL puromycin (Invitrogen, ant-pr-1) for selection. Following an additional 72 hr of culture, viable cell recovery was assessed by flow cytometry using eFluor 506 viability dye (Thermo Fisher, 65-0866-18).

Functional viral titre was determined using the following equation, where the percentage of live cells was normalized to the percentage of live cells in the corresponding non-puromycin-treated control:

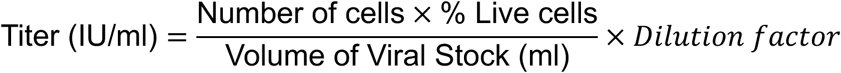

The volume of lentivirus required to achieve an MOI of 0.3 was calculated using the following formula:

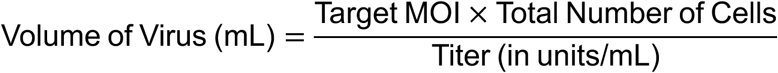

### Pooled CRISPR-Cas9 knockout screen

The pooled CRISPR-Cas9 knockout T cells were manufactured using a modified single guide RNA (sgRNA) lentiviral infection with Cas9 protein electroporation (SLICE) approach, as previously described by Shifrut et al (*28*). Primary human CD3⁺ T cells were activated on day 0 and transduced with the pooled lentiviral sgRNA library (GenScript) on day 1 at a multiplicity of infection (MOI) of 0.3 to maximize the likelihood of a single sgRNA integration per cell. On day 2, transduced T cells were electroporated with Cas9 (IDT, 1081059; knockout condition) or no Cas9 (library transduced condition). Prior to electroporation, cells were resuspended in 100μL per 10×10^6^ cells of Lonza P3 Primary Cell Nucleofector Solution. Per 100 µL Nucleocuvette® (Lonza, V4XP-4024), 30μg of recombinant Cas9 protein (IDT, 1081059) was electroporated with 10×10^6^ T cells using the Lonza 4D-Nucleofector at pulse code EO-115. Immediately following electroporation, 1mL of pre-warmed ImmunoCult™-XF T Cell Expansion Medium (no cytokines) was added to each cuvette and incubated for 20 mins at 37°C before cells were returned to culture. Transduced cells were cultured in ImmunoCult™-XF T Cell Expansion Medium and selected with puromycin (Invitrogen, ant-pr-1, 2μg/mL) from days 4–12 to enrich for sgRNA-expressing cells. On day 12, cells were transferred to 100% HGSOC ascites supernatant (Patient 3, Fig. 1I-J) and reactivated. For Supplementary Fig. 1E, on day 14, bulk knockout T cell or library-transduced T cell phenotypes were assessed by flow cytometry, measuring IFN-γ and mitochondrial membrane potential. For Fig. 3A-B, cells were sorted using a FACSAria Fusion and BD FACSAria III cell sorters into IFN-γ-high /ΔΨm-high and IFN-γ-low/ΔΨm-low populations. Sorted cells were collected directly into PBS before genomic DNA extraction.

Screen coverage was maintained throughout the experiment by expanding sufficient numbers of cells to preserve at least 100-fold representation of each sgRNA, consistent with established recommendations for pooled CRISPR screening. Coverage calculations incorporated the total library size (29,903 sgRNAs), anticipated cell expansion (3-fold), and expected fluorescence-activated cell sorting (FACS) recovery to ensure adequate representation of all library elements in downstream analyses.

### sgRNA library design, preparation, and next-generation sequencing

A custom metabolism-focused sgRNA library comprising 29,903 sgRNAs targeting 2,981 metabolic genes, cloned into a lentiviral expression vector, was generated by GenScript. The library gene list is based on a previously published human CRISPR knockout library, described in Birsoy et al (*61*). Library synthesis, cloning into the pLentiGuide Puro vector, plasmid amplification, and quality control were performed by GenScript (Piscataway, NJ, USA).

Following FACS, genomic DNA was isolated from the sorted high-effector function (IFN-γ-high /ΔΨm-high) and low-effector function (IFN-γ-low/ΔΨm-low) populations using a phenol-chloroform extraction protocol. Briefly, cells were lysed in SDS-containing lysis buffer (1% SDS, 50 mM Tris, pH 8, 10 mM EDTA) followed by RNase A (Zymo, E1008) and proteinase K (Zymo, D3001) digestion. Genomic DNA was purified by phenol:chloroform alcohol extraction using Phase Lock Gel tubes (Quantabio, 2302820), precipitated with sodium acetate (3M) and isopropanol, washed with 70% ethanol, and resuspended in UltraPure DNase/RNase-Free Distilled Water (Thermo Fisher, 10977015).

Integrated sgRNA sequences were amplified by the McGill University and Genome Québec Innovation Centre (Montréal, QC, Canada) using primers flanking the integrated sgRNA cassette. Amplicons were subjected to next-generation sequencing on an Illumina NextSeq platform according to the Genome Québec workflow. Sequencing reads were demultiplexed and processed for downstream analysis.

A list of genes included in the library is shown in Table S4.

### CRISPR screen analysis

Sequencing reads were processed and quantified using the Model-based Analysis of Genome-wide CRISPR-Cas9 Knockout (MAGeCK) software. sgRNA read counts from the input library and the sorted high-effector function (IFN-γ-high /ΔΨm-high) and low-effector function (IFN-γ-low/ΔΨm-low populations) were normalized to control (intergenic) sgRNAs prior to analysis.

Gene-level enrichment was assessed using the MAGeCK maximum likelihood estimation (MLE) algorithm by comparing sgRNA abundance between the high-effector function and low-effector function populations (*30*, *62*). Positive enrichment scores indicate genes whose disruption enhanced T cell function under suppressive ascites conditions, whereas negative enrichment scores indicate genes depleted from the high-effector function population. Candidate genes were ranked according to MAGeCK enrichment Z-scores and associated P values.

Sequencing quality was evaluated by determining the percentage of reads successfully mapped to sgRNAs within the custom CRISPR library and by calculating the Gini index to assess sgRNA representation across samples (MAGeCK-VISPR). Principal component analysis (PCA) of normalized sgRNA counts was performed to evaluate the separation of the input library and sorted populations.

Data visualization and statistical analyses were performed using R (version 4.4.2), MAGeCK-VISPR and GraphPad Prism 11.0.1.

Data from pooled CRISPR-Cas9 screen is presented in Table S5.

### CRISPR RNP-mediated gene editing

Individual gene knockouts were generated by electroporation of Cas9 ribonucleoprotein (RNP) complexes into activated primary human T cells. Chemically synthesized sgRNAs targeting peroxiredoxin 1-4 (PRDX1-4), or control loci (AAVS1 and intergenic) were obtained from IDT.

Cas9 RNP complexes were assembled by combining 100 pmol sgRNA with 48.8 pmol recombinant Cas9 protein (IDT, 1081059), corresponding to an approximate 2:1 sgRNA:Cas9 molar ratio, and incubating at 37°C for 15 mins. Activated T cells were electroporated at 10⁶ cells per condition using the P3 Primary Cell 4D-Nucleofector X Kit S (Lonza, V4XP-3032) in 20-µL Nucleocuvette™ Strip wells (Lonza). Electroporation was performed using a Lonza 4D-Nucleofector™ with program EO-115. Immediately following electroporation, 80 µL of pre-warmed ImmunoCult™-XF T Cell Expansion Medium (STEMCELL Technologies, 10981; no cytokines) was added to each cuvette well and incubated for 15 mins at 37°C before cells were returned to culture. Gene editing efficiency was either assessed 4 or 11 days after electroporation by Tracking of Indels by Decomposition (TIDE) analysis of genomic DNA amplicons and/or by western blot analysis of PRDX1 protein expression.

All sgRNA sequences used for single knockout experiments can be found in Table S3.

### TIDE analysis

Gene editing efficiency was assessed by TIDE analysis (*63*). Genomic DNA was isolated from edited T cells 4 or 11 days following CRISPR-Cas9 RNP electroporation (7 or 14 days after T cell activation) using DNA QuickExtract (Lucigen, QE09050), according to manufacturer’s instructions. The genomic region surrounding each sgRNA target site was amplified by PCR using gene-specific primers (designed using CRISPOR web tool (*64*), https://crispor.gi.ucsc.edu/) and subjected to Sanger sequencing at the Genomics Center, CHU de Québec–Université Laval Research Center. PCR was performed using Phusion High-Fidelity PCR Master Mix (Thermo Fisher, F531L), with reaction mixtures and thermal cycler conditions carried out per manufacturer’s instructions.

Sanger sequencing chromatograms from edited and control samples were analyzed using the TIDE web tool (https://tide.nki.nl/) to estimate insertion and deletion (indel) frequencies generated by CRISPR-Cas9-mediated gene editing. Editing efficiency was calculated using default analysis parameters.

All primer sequences can be found in Table S3.

### Western blotting

Protein lysates were prepared from an equal number of cells per condition. Cells were pelleted by centrifugation and lysed in RIPA buffer (Thermo Fisher, 89900) supplemented with Halt Protease and Phosphatase Inhibitor Cocktail (Thermo Fisher, 87786). Lysates were incubated on ice for 30 min and clarified by centrifugation at 13,000 rpm for 10 min at 4°C. Supernatants were collected and stored at −80°C or used immediately for downstream analysis. Protein concentration was determined using the Pierce BCA Protein Assay (Thermo Fisher, 23225).

Samples were prepared in NuPAGE™ LDS Sample Buffer (Thermo Fisher, NP0007) and NuPAGE™ Sample Reducing Agent (Thermo Fisher, NP0009), heated at 70°C for 10 min, and resolved on NuPAGE™ 4–12% Bis-Tris gels (Thermo Fisher, NP0336BOX) using MES running buffer. Gels were run for 60–75 min at 125 V using an Invitrogen Mini Gel Tank apparatus. Proteins were transferred to nitrocellulose membranes (Thermo Fisher, LC2006) using semi-dry transfer conditions for 45 min at 22 V using Invitrogen Power Blotter.

Membranes were blocked for 1 hr in Odyssey Blocking Buffer (LI-COR Biosciences, 927-60001) and incubated overnight at 4°C with primary antibodies diluted in blocking buffer. Membranes were washed in TBST and incubated for 1hr at room temperature with infrared dye-conjugated secondary antibodies. Blots were washed and imaged using an Odyssey M Imaging System. Protein abundance was quantified by densitometry and normalized to β-actin.

All western blot antibodies can be found in Table S2.

### Reverse Transcription quantitative PCR (RT-qPCR)

Total RNA was extracted from primary human T cells using the RNeasy Plus Mini Kit (QIAGEN, 74134) according to the manufacturer’s instructions. RNA concentration was determined by NanoDrop spectrophotometry, and purified RNA was stored at −20°C until further use. Complementary DNA (cDNA) was synthesized from 500 ng of total RNA using the qScript cDNA Synthesis Kit (QuantaBio, 95047). Quantitative PCR was performed using TaqMan Fast Advanced Master Mix (Thermo Fisher, 4444557) and 20× TaqMan Gene Expression Assays (Thermo Fisher; Hs00602020_mH for PRDX1, Hs02786624_g1 for GAPDH [used for normalization]), according to manufacturers instructions. Reactions were run using the following cycling conditions: 50°C for 2 min, 95°C for 2 min, followed by 40 cycles of 95°C for 1 s and 60°C for 20 s. Each sample was analyzed in four technical replicates and included no-template and no-reverse-transcriptase controls. Relative PRDX1 expression was normalized to GAPDH and calculated using the 2^-ΔCt^ method.

### CAR-T cell generation

Primary human CD3⁺ T cells were isolated from PBMCs obtained from healthy donor leukopaks (STEMCELL Technologies) using Miltenyi CD3 MicroBeads (130-050-101, per manufacturers protocol) and activated on day 0 with anti-CD3/CD28 T cell activators (STEMCELL Technologies, 10971) at 25 µL per 10⁶ cells. Cells were cultured at 10⁶ cells/mL in ImmunoCult-XF T Cell Expansion Medium (STEMCELL Technologies, 10981) supplemented with 1% penicillin-streptomycin and recombinant human IL-2 (PeproTech, 200-02-100UG).

On day 1, activated T cells were transduced with lentivirus encoding a folate receptor α (FRα)-specific chimeric antigen receptor (CAR) and a green fluorescent protein (GFP) reporter (GenScript). The CAR construct sequence was obtained from a University of Pennsylvania patent (*36*, *65*). Cells were transduced with 3µL (multiplicity of infection [MOI] unknown) of concentrated lentiviral supernatant. CAR-transduced cells were split on day 3 and enriched on day 4 by magnetic selection. Briefly, cells were stained with an anti-G4S linker antibody (Cell Signaling Technologies, 69782L) for 20 min at room temperature in the dark, followed by labeling with anti-AF647 MACS microbeads (Miltenyi, 130-091-395) and magnetic enrichment using Miltenyi LS columns according to the manufacturer’s instructions. Enriched CAR-T cells were returned to culture at 2×10^5^ cells/mL in ImmunoCult-XF T Cell Expansion Medium (STEMCELL Technologies, 10981).

On day 6, CAR-T cells were subjected to CRISPR-Cas9 RNP-mediated gene editing. RNP complexes were prepared using recombinant Cas9 protein and sgRNAs targeting PRDX1 or control conditions, as indicated. Cells were resuspended in Lonza P3 Nucleofector Solution (V4XP-3032) at 20 µL per 10⁶ cells and mixed with preassembled RNP complexes before transfer to Lonza 20-µL 16-well Nucleocuvette strips (V4XP-3032). Electroporation was performed using the Lonza 4D-Nucleofector with pulse code EO-115. See section “*CRISPR RNP-mediated gene editing”* for a more detailed protocol.

CAR expression was assessed by flow cytometry on day 12 based on GFP positivity, and CAR-positive cell numbers were used to normalize effector cell input for downstream cytotoxicity assays. Edited CAR-T cells were either used in cytotoxicity assays or cultured in complete medium or patient HGSOC ascites with anti-CD3/CD28/CD2 restimulation (STEMCELL Technologies) for 48 hrs prior to flow cytometric analysis.

### CAR-T cell cytotoxicity assay

Cytotoxic activity was assessed using luciferase-expressing SKOV3 ovarian cancer cells (SKOV3-Luc). SKOV3-Luc cells were maintained in McCoy’s 5A medium (Gibco, 16600082) supplemented with 10% fetal bovine serum (FBS) and 1% penicillin-streptomycin. One day prior to co-culture, SKOV3-Luc cells were plated at 2×10⁴ cells per well in white flat-bottom 96-well plates and allowed to adhere overnight. On the day of the assay, patient HGSOC ascites supernatant was thawed, filtered, and added to the appropriate wells, while media wells received complete RPMI medium (Cytivia, SH30027.01; supplemented with 5% human serum, 12.5 mM HEPES, 2mM L-glutamine, and 1% pen-strep). CAR-T cells were harvested, washed, and resuspended in either complete RPMI or ascites at 2×10⁵ CAR⁺ cells/mL, where CAR⁺ cell numbers were calculated by multiplying the total viable T cell count by the percentage of GFP-positive cells determined by flow cytometry. CAR-T cells were added to SKOV3-Luc cells at an effector-to-target (E:T) ratio of 10:1, 5:1, 2.5:1, 1.25:1, 1:1, 0.625:1 or 0.3125:1 and co-cultured for 24 h. Tumor-only and media-only wells were included as controls.

Following co-culture, D-luciferin (Revvity, 122799) was diluted to a 10x working solution in PBS and added to each well to achieve a final 1x concentration. Plates were incubated for 5 min at room temperature in the dark before luminescence was measured using a plate reader. Tumor cell killing was quantified as percent specific killing using the following equation:

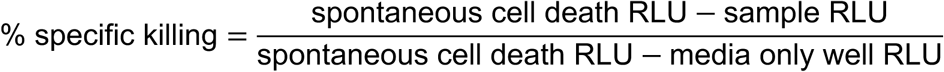

where RLU represents relative luminescence units.

### Autophagic flux assay

Autophagic flux was assessed in primary human T cells by measuring LC3-II accumulation following pharmacological inhibition (using hydroxychloroquine [HCQ]) of lysosomal degradation. *PRDX1*-knockout (treated with *PRDX1-*targeting sgRNA 10) and control T cells (treated with a sgRNA targeting an intergenic region) were cultured under standard expansion conditions and treated on day 13 following initial activation (one day following restimulation). Cells were left untreated or treated with rapamycin (Cayman Chemical, 13346-10; autophagy stimulator), HCQ (Sigma, H0915-5MG; autophagy inhibitor) or a combination of rapamycin and HCQ for 20 h before protein lysate collection. Rapamycin (Cayman Chemical, 13346-10) was added to a final concentration of 400 nM, and HCQ (Sigma, H0915-5MG) was added to a final concentration of 30 µM.

Following treatment, whole-cell lysates were collected on day 14 and analyzed by western blotting for LC3, as described below. Equal numbers of cells were collected for each condition, and equivalent lysate volumes were loaded for electrophoresis. β-actin served as the loading control.

### Oxidative stress assay

To assess susceptibility to oxidative stress, primary human T cells (control or *PRDX1-*knockout T cells) were restimulated 12 days after isolation/initial activation with anti-CD3/CD28/CD2 T cell activators (STEMCELL Technologies, 10970) and cultured in the presence or absence of hydrogen peroxide (H₂O₂). Cells were plated at 10⁶ cells/mL and treated with 100 µM H₂O₂ for 48 hrs for H₂O₂ treated condition. Control samples consisted of a pooled mixture of cells transfected with no sgRNA, an *AAVS1*-targeting sgRNA, or an intergenic sgRNA, whereas *PRDX1*-knockout samples consisted of a pooled mixture of cells transfected with *PRDX1*-targeting sgRNAs 1, 8, and 10 (Fig. 3).

Following incubation, cell viability was assessed by flow cytometry using eFluor 506 Fixable Viability Dye (Thermo Fisher, 65-0866-18) according to the manufacturer’s instructions. Flow cytometry acquisition and analysis were performed as described above.

### Statistical analysis

Statistical analyses were performed using GraphPad Prism (version 11.1). The statistical tests used for each experiment are indicated in the corresponding figure legends. Unless otherwise indicated, each technical replicate (represented by data points) represents an independent experiment or conditions within the same experiment that were targeted with a unique sgRNA (e.g. no sgRNA, intergenic sgRNA or AAVS1 sgRNA in control condition; PRDX1 sgRNA 1, 8, 10 for PRDX1 knockout condition). Biological replicates (e.g. distinct patient ascites or T cell donors) are represented by different plots, or different conditions on the same plot.

For comparisons between two groups, two-tailed Welch’s t-tests or paired t-tests were performed, as appropriate. P values < 0.05 were considered statistically significant. For PCA analysis, group separation was determined as statistically significant by PERMANOVA analysis. For pooled CRISPR-Cas9 screening data, gene-level enrichment was determined using the MLE algorithm implemented in MAGeCK. Candidate genes were ranked according to enrichment Z-scores and associated P values.

## Acknowledgments

We would like to thank members of the Terry Fox Labs Flow Cytometry Core, including Alynn Shanks, Guillermo Simkin, Wenbo Xu, and Christopher May, who heavily assisted in the fluorescently activated cell sorting. We also thank GenScript for the custom sgRNA library generation. Furthermore, we thank Genome Quebec for their services in next generation sequencing for the pooled CRISPR-Cas9 screen. We thank members of the BC Cancer Tumor Tissue Repository (TTR), including Dr. Peter Watson, Tamsin Tarling, Sindy, Simon Dee, Jodi LeBlanc and Tim Shack who provide invaluable services in managing patient samples. Lastly, thank you to Dr. Julian Smazynski and his team for gifting us the luciferase-expressing SKOV3 cells used in this study.

## Funding

This work was supported by the Terry Fox Research Institute New Frontiers Program Project Grant (MetaboHUB; Project No. 1125) and the Canadian Institutes of Health Research (CIHR; PJT-192015). S.J.M. was supported by a Cancer Research Society Doctoral Research Award and a BC Cancer Rising Star Award.

## Author contributions

J.J.L., G.C. and S.J.M conceived the study. S.J.M designed and performed the experiments, interpreted the results, and wrote the manuscript. G.C. and S.M. performed experiments. S.P. and L.C. assisted with experiments. S.H performed the pooled CRISPR-Cas9 screen analysis. S.J.M and J.J.L edited the manuscript. All authors revised the manuscript.

## Competing interests

The authors declare no competing interests.

## Data and materials availability

Pooled CRISPR-Cas9 screen data is provided in Table S5. Antibodies and primer sequences are provided in Tables S2-S3, respectively.

## List of Supplementary Figures

**Supplementary Fig. 1.**
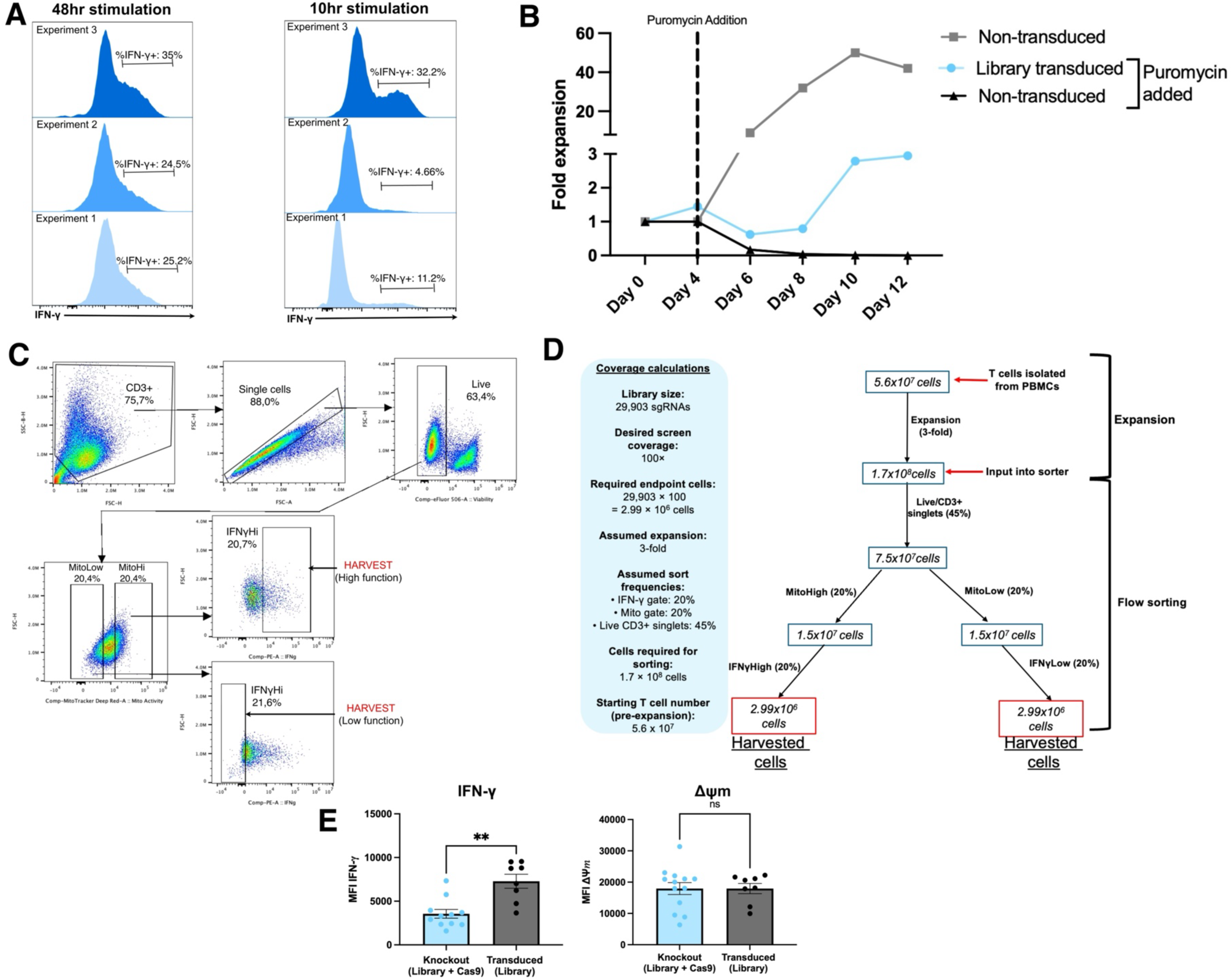
Optimization of the pooled CRISPR-Cas9 screening workflow. **(A)** Optimization of intracellular IFN-γ detection for pooled CRISPR screening. Representative histograms showing IFN-γ expression following 10 or 48 hr stimulation. Experiments were performed using T cells from the same donor under standard media conditions. **(B)** CD3+ T cells were transduced with the pooled sgRNA library and subsequently subjected to puromycin. Fold expansion of non-transduced cells cultured without puromycin, library-transduced cells cultured with puromycin, and non-transduced cells cultured with puromycin is shown. **(C)** Complete flow cytometry gating strategy used for pooled CRISPR screening. **D)** Coverage calculations used to guide pooled CRISPR screen design. Based on a library containing 29,903 sgRNAs and a desired coverage of 100 cells per sgRNA, calculations were performed to estimate the number of cells required at each stage of the workflow to ensure adequate representation of all sgRNAs in the sorted high-effector function and low-effector function populations. **(E)** Functional comparison of pooled knockout and non-knockout library populations. T cells transduced with the pooled sgRNA library and electroporated with Cas9 protein (pooled knockout population) were compared with cells transduced with the pooled sgRNA library alone (non-knockout population). Intracellular IFN-γ expression and mitochondrial membrane potential were assessed following a 48 hr stimulation. Data are shown from a single donor; each point represents a technical replicate obtained within or across experiments. Bars represent mean ± SEM. Statistical significance was determined using Welch’s t-test. **P < 0.01; ns, not significant.

**Supplementary Fig. 2.**
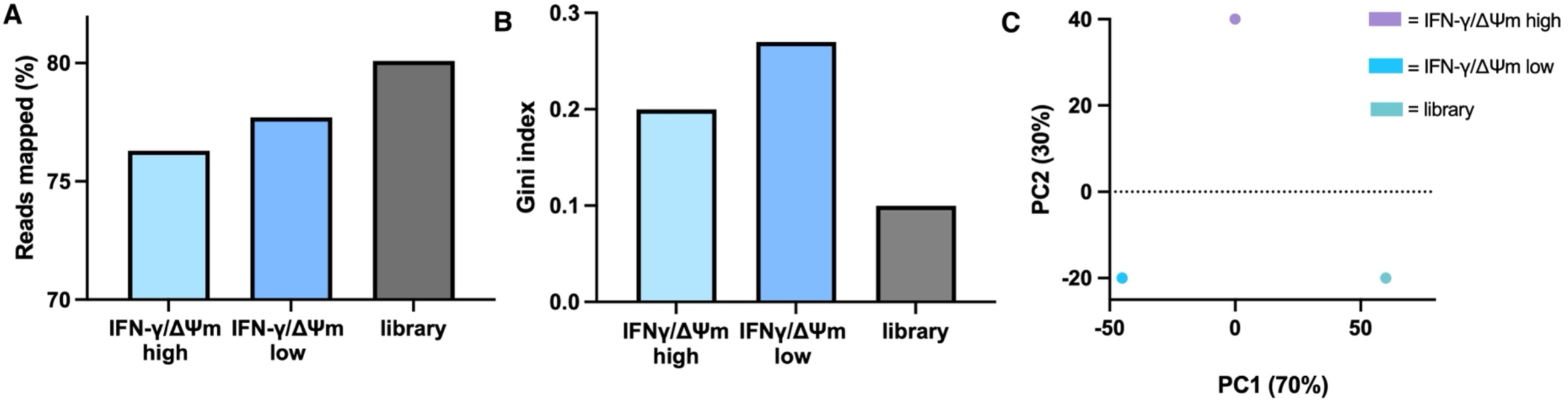
Sequencing quality-control analyses demonstrate adequate sgRNA representation and separation of sorted screening populations. **(A)** Percentage of sequencing reads successfully mapped to sgRNAs contained within the pooled CRISPR library. Mapping efficiency is shown for the input library, high-effector function (IFN-γ-high, ΔΨm-high) population, and low-effector function (IFN-γ-low, ΔΨm-low) population recovered from the pooled CRISPR screen. Sequencing was performed using an Illumina NextSeq platform. **(B)** Gini index analysis of sgRNA abundance distributions within the input library and sorted populations. **(C)** Principal component analysis (PCA) of MAGeCK-normalized sgRNA read counts from the input library and sorted populations. Data were generated from the same pooled CRISPR screen shown in Fig. 3.

**Supplementary Fig. 3.**
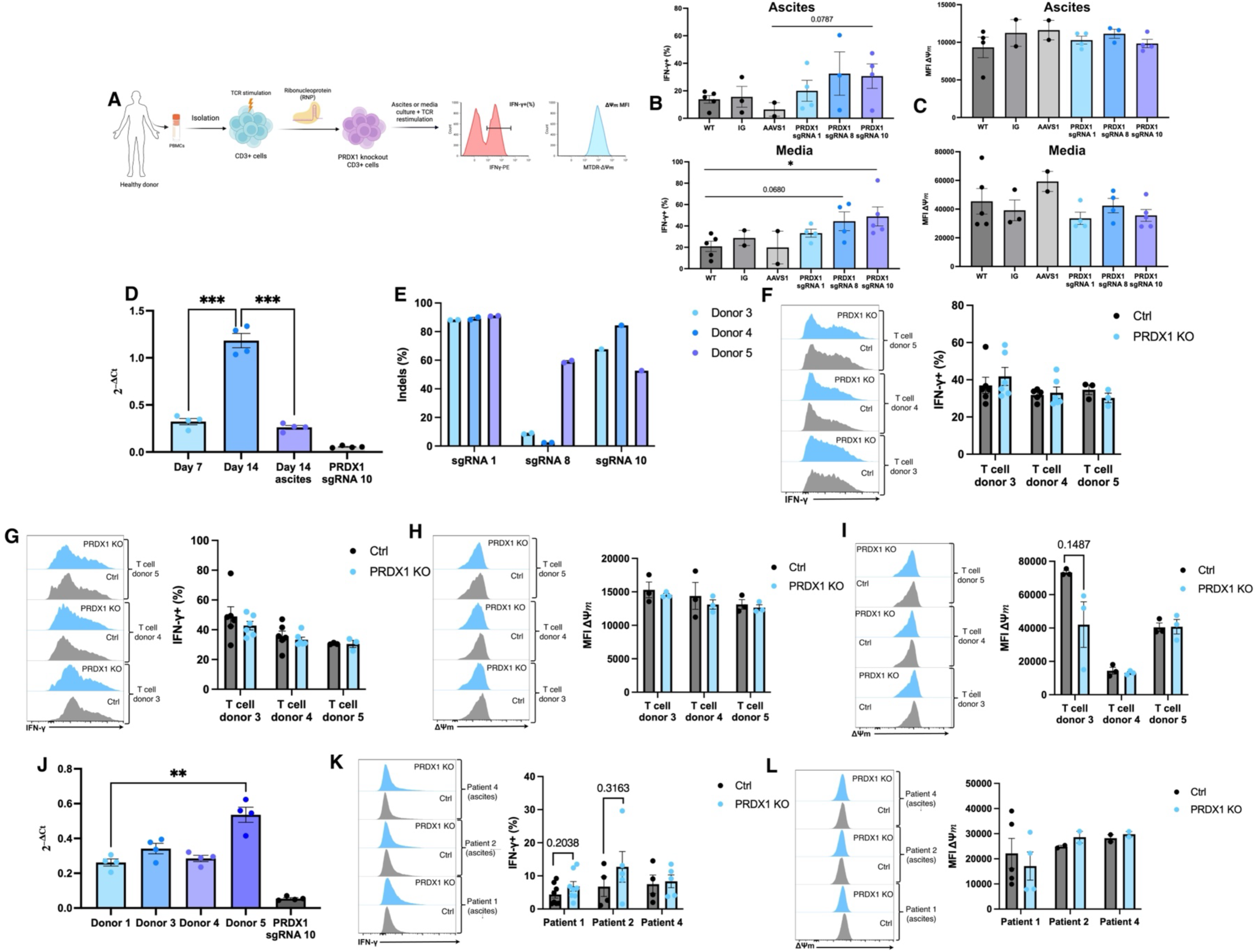
The effects of *PRDX1* deletion on IFN-γ production are reproducible across multiple sgRNAs but vary across donors and ascites samples. **(A)** Experimental workflow used for validation of *PRDX1* deletion. CD3+ T cells were isolated from healthy donors, activated, electroporated with *PRDX1*-targeting ribonucleoprotein (RNP) complexes, cultured in media or ascites, and restimulated prior to assessment of intracellular IFN-γ production and mitochondrial membrane potential (ΔΨm). Created with BioRender. **(B)** Validation of independent *PRDX1*-targeting sgRNAs. IFN-γ production was measured following 48 hr stimulation in either ascites or media. Data are shown from the same donor used in the pooled CRISPR screen (Fig. 3). Each point represents a technical replicate performed within the same experiment or across multiple experiments. **(C)** Mitochondrial membrane potential (ΔΨm) following treatment with independent *PRDX1-*targeting sgRNAs. Data are shown from the same donor and experimental conditions described in panel (B). **(D)** Relative PRDX1 mRNA expression (2^-ΔCt^), normalized to GAPDH, in primary human T cells at day 7, day 14 in media, and day 14 in HGSOC ascites supernatant. In both day 14 samples, T cells were restimulated 48 hrs prior. Each point represents an individual technical replicate. **(E)** Indel generation efficiency of *PRDX1*-targeting sgRNAs identified in the CRISPR screen. Indel frequencies were determined by TIDE analysis of genomic DNA harvested from expanded T cells 14 days after isolation and activation and 11 days following Cas9 ribonucleoprotein (RNP) electroporation. Data were generated for sgRNA 1, 8, and 10 (Fig. 3B) in T cell donors 3–5. **(F–G)** IFN-γ production following *PRDX1* deletion in additional T cell donors. Representative histograms and quantification of intracellular IFN-γ-positive cells are shown for donor 3, donor 4, and donor 5 following 48 hr stimulation in ascites **(F**) or media **(G). (H–I)** Mitochondrial membrane potential (ΔΨm) following *PRDX1* deletion in additional T cell donors. Representative histograms and quantification of MT-DR mean fluorescence intensity (MFI) are shown for donor 3, donor 4, and donor 5 following 48 hr stimulation in ascites **(H)** or media **(I**). **(J)** Relative PRDX1 mRNA expression (2^-ΔCt^), normalized to GAPDH, in stimulated T cells cultured in HGSOC ascites supernatant from healthy donors 1, 3, 4, and 5, together with *PRDX1*-knockout T cells generated using sgRNA 10. **(K)** IFN-γ production following *PRDX1* deletion in ascites supernatant obtained from three independent HGSOC patients. Representative histograms and quantification of intracellular IFN-γ-positive cells are shown following 48 hr stimulation. Experiments were performed using the same donor used in the pooled CRISPR screen and primary validation studies (Figs. 3 and 4). **(L)** Mitochondrial membrane potential (ΔΨm) following *PRDX1* deletion in ascites supernatant obtained from three independent HGSOC patients. Representative histograms and quantification are shown using the same conditions described in panel (K). For all panels, statistical significance was determined using Welch’s t-test. Control conditions included wild-type, intergenic sgRNA, or AAVS1-targeting sgRNA controls, whereas PRDX1-knockout cells were generated using PRDX1-targeting sgRNAs (sgRNA 1, 8 or 10). Bars represent mean ± SEM. *P < 0.05, **P < 0.01.

**Supplementary Fig. 4.**
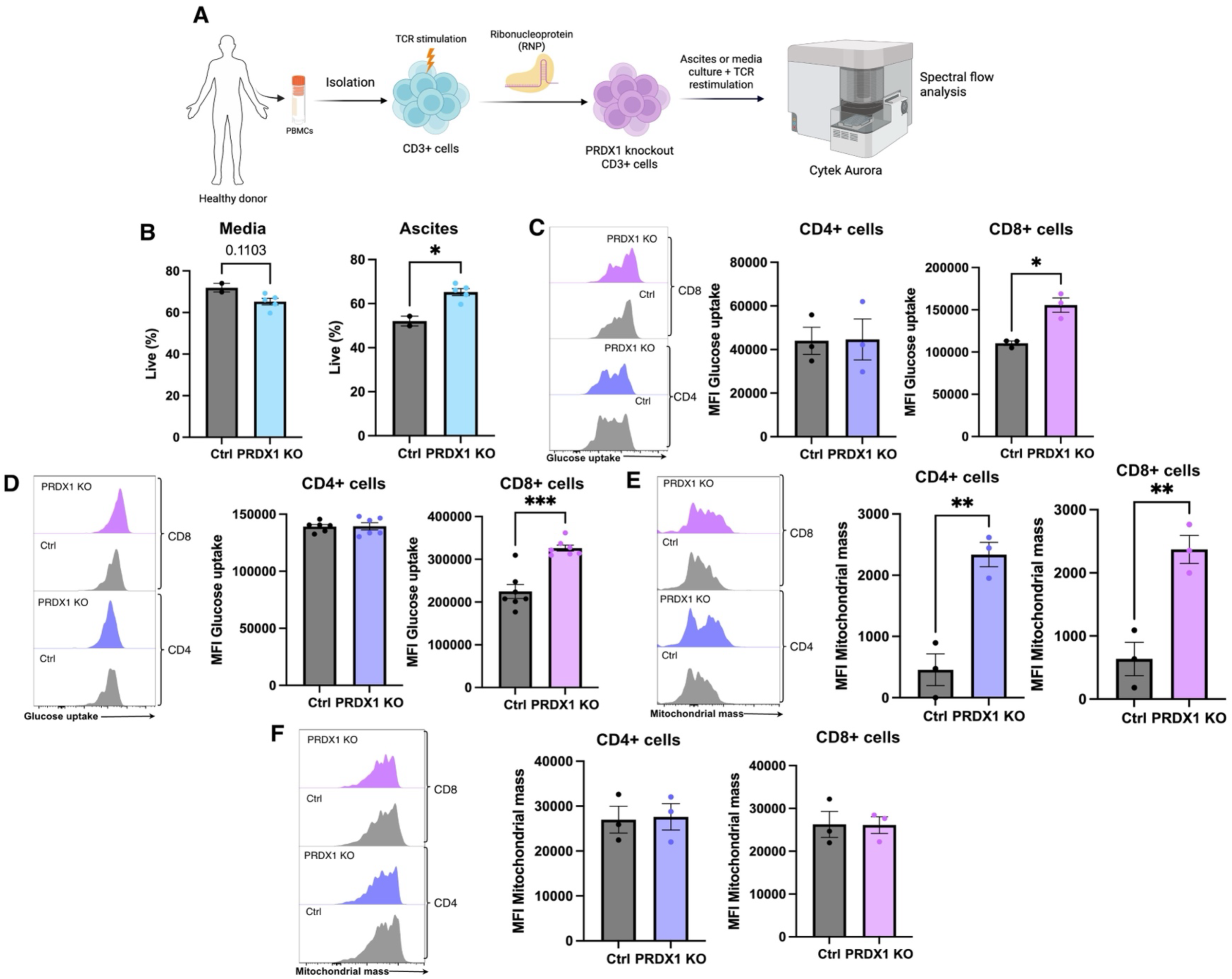
*PRDX1* deletion increases glucose uptake and mitochondrial mass in ascites-cultured T cells. **(A)** Experimental workflow used to assess metabolic and functional phenotypes following *PRDX1* deletion. CD3+ T cells were isolated from healthy donors, activated, electroporated with *PRDX1*-targeting ribonucleoprotein (RNP) complexes, cultured in media or ascites, restimulated with anti-CD3/CD28/CD2 antibodies, and analyzed by spectral flow cytometry. Created with BioRender. **(B)** Viability of control and *PRDX1*-knockout T cells following 48 hr stimulation in media or HGSOC ascites. **(C)** Glucose uptake in ascites-cultured T cells. Representative histograms and mean fluorescence intensity of GlucoseCy5 uptake are shown for CD4+ and CD8+ T cell subsets following 48 hr stimulation of CD3+ T cells in HGSOC ascites. **(D)** Glucose uptake in media-cultured T cells. Representative histograms and mean fluorescence intensity of GlucoseCy5 uptake are shown for CD4+ and CD8+ T cell subsets following 48 hr stimulation of CD3+ T cells in media. **(E)** Mitochondrial mass in ascites-cultured T cells. Representative histograms and quantification of mitochondrial mass are shown for CD4+ and CD8+ T cell subsets following 48 hr stimulation of CD3+ T cells in HGSOC ascites. **(F)** Mitochondrial mass in media-cultured T cells. Representative histograms and quantification of mitochondrial mass measurements are shown for CD4+ and CD8+ T cell subsets following 48 hr stimulation of CD3+ T cells in media. For panels B–F, each data point represents a distinct sgRNA tested in one donor sample. Control samples included no sgRNA, intergenic sgRNA, or *AAVS1*-targeting sgRNA, whereas *PRDX1*-knockout samples included PRDX1 sgRNA1, sgRNA8, or sgRNA10. Some sgRNAs were evaluated in technical replicates across multiple experiments. T cells were cultured in media or ascites derived from the same HGSOC patient sample used in the pooled CRISPR screen (Fig. 1I-J: patient 3). Bars represent mean ± SEM. Statistical significance was determined using Welch’s t-test. *P < 0.05, **P < 0.01, ***P < 0.001.

**Supplementary Fig. 5.**
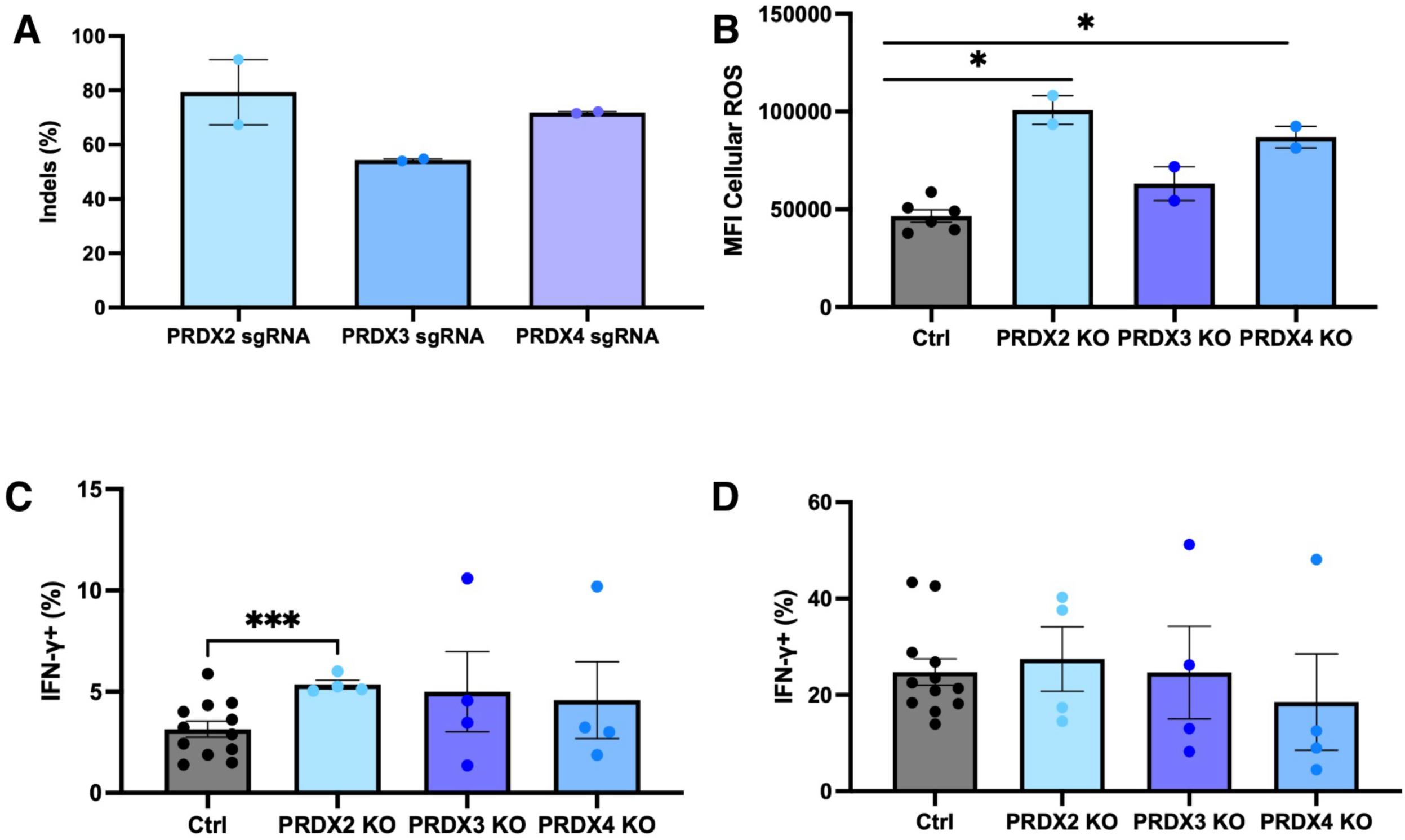
PRDX family members differentially regulate T cell ROS and effector function. **(A)** Validation of CRISPR-mediated deletion of *PRDX2*, *PRDX3*, and *PRDX4* genes. Gene editing efficiency was determined by TIDE analysis of genomic DNA harvested on day 14 of T cell expansion (11 days following RNP electroporation). **(B)** Cellular ROS following deletion of *PRDX2*, *PRDX3*, or *PRDX4*. Cellular ROS was measured following 48 hr stimulation in media. **(C)** Percentage of IFN-γ producing CD3+ cells following deletion of *PRDX2*, *PRDX3*, or *PRDX4* in HGSOC ascites (ascites derived from the same HGSOC patient sample used in the pooled CRISPR screen [Fig. 1I-J: patient 3]). Intracellular IFN-γ was measured following 48 hr stimulation. **(D)** Percentage of IFN-γ producing CD3+ cells following deletion of *PRDX2*, *PRDX3*, or *PRDX4* under media conditions. Intracellular IFN-γ was measured following 48 hr stimulation. For A-D, data are shown for one T cell donor, with each point representing a technical replicate performed within or across experiments. Bars represent mean ± SEM. Statistical significance was determined using Welch’s t-test. Control samples included no sgRNA, intergenic sgRNA, or *AAVS1-*targeting sgRNA controls. *P < 0.05, ***P < 0.001.

## List of Supplementary Tables

Table S1: Patient information

Table S2: Antibodies

Table S3: Primer and sgRNA sequences

Table S4: CRISPR-Cas9 library gene list

Table S5: CRISPR screen data (MLE gene summary)

## Notes

### Competing Interest Statement

The authors have declared no competing interest.

